# Adhesion-based capture stabilizes nascent microvilli at epithelial cell junctions

**DOI:** 10.1101/2023.03.08.531705

**Authors:** Caroline S. Cencer, Jennifer B. Silverman, Leslie M. Meenderink, Evan S. Krystofiak, Bryan A. Millis, Matthew J. Tyska

**Affiliations:** Department of Cell and Developmental Biology, Vanderbilt University School of Medicine, Nashville, TN 37232; Department of Biomedical Engineering, Vanderbilt University School of Engineering, Nashville, TN 37235

**Keywords:** transporting epithelia, actin, cadherins, differentiation, brush border, intermicrovillar adhesion

## Abstract

Differentiated transporting epithelial cells present an extensive apical array of microvilli – a “brush border” – where neighboring microvilli are linked together by intermicrovillar adhesion complexes (IMACs) composed of protocadherins CDHR2 and CDHR5. Although loss-of-function studies provide strong evidence that IMAC function is needed to build a mature brush border, how the IMAC contributes to the stabilization and accumulation of nascent microvilli remains unclear. We found that, early in differentiation, the apical surface exhibits a marginal accumulation of microvilli, characterized by higher packing density relative to medial regions of the surface. While medial microvilli are highly dynamic and sample multiple orientations over time, marginal protrusions exhibit constrained motion and maintain a vertical orientation. Unexpectedly, we found that marginal microvilli span the junctional space and contact protrusions on neighboring cells, mediated by complexes of CDHR2/CDHR5. FRAP analysis indicated that these *transjunctional* IMACs are highly stable relative to adhesion complexes between medial microvilli, which explains the restricted motion of protrusions in the marginal zone. Finally, long-term live imaging revealed that the accumulation of microvilli at cell margins consistently leads to accumulation in medial regions of the cell. Collectively, our findings suggest that nascent microvilli are stabilized by a capture mechanism that is localized to cell margins and enabled by the transjunctional formation of IMACs. These results inform our understanding of how apical specializations are assembled in diverse epithelial systems.

## INTRODUCTION

Organ function depends on specialized cell types that have evolved morphologies to enable specific physiological tasks. Transporting epithelial cells, found in the intestine and kidney proximal tubule, offer interesting examples of this phenomenon. As important sites of solute uptake, maximizing apical surface area is a critical facet of these tissues. To meet this challenge, epithelial cells extend 1000s of bristle-like protrusions called microvilli, which pack tightly into a highly ordered array to collectively form a brush border [1, 2]. A single microvillus is a cylinder-shaped, micron-scale membrane protrusion supported by a core actin bundle consisting of 20-40 actin filaments [3, 4]. By scaffolding apical membrane in this way, microvilli amplify surface area available for solute transport and optimize solute uptake potential [5–7]. Microvilli first appear on the cell surface very early in epithelial maturation; differentiating cells, like those found within intestinal stem cell-containing crypts, exhibit few, poorly organized microvilli [8]. However, differentiated, fully functional enterocytes, found on the villus or within the kidney tubule, present a well-organized brush border consisting of densely packed protrusions [3, 9, 10].

Several previous studies established that tight microvillar packing is driven by a protocadherin-based intermicrovillar adhesion complex (IMAC), which physically links the distal tip of a microvillus to the tips of its neighboring microvilli [11–14]. In the case of the enterocyte, these adhesive interactions give rise to a hexagonal packing pattern when viewed *en face*, which represents maximum surface occupancy. Previous work also identified protocadherins CDHR2 and CDHR5 as the primary adhesive elements in these links, which form *trans* heterophilic adhesion complexes that are well suited for bridging the ∼50 nm gap between neighboring microvilli [11, 15]. CDHR2 and CDHR5 ectodomains contain multiple extracellular cadherin (EC) repeat motifs arranged in tandem, which are anchored to the membrane via a single spanning transmembrane domain [16]. Both protocadherins also contain cytoplasmic tails at their C-termini, which enable direct interactions with cytoplasmic IMAC binding partners including the actin-based motor, myosin-7B (MYO7B), and the scaffolding proteins, ankyrin repeat and sterile alpha motif domain containing 4B (ANKS4B) and usher syndrome 1C (USH1C) [12, 13, 17, 18]. Recently, calmodulin-like protein 4 (CALML4) was identified as a direct binding partner of MYO7B, making it an additional IMAC component [19]. KD studies of MYO7B indicate that this motor plays a key role in the localization of CDHR2/CDHR5 adhesion complexes to the distal tips of microvilli [12, 13]. In the differentiating CACO-2_BBE_ intestinal epithelial cell culture model, disrupting the function of the IMAC via calcium chelators or knockdown of any single complex component leads to striking defects in microvillar growth and packing organization during differentiation [11–13, 19]. Furthermore, complete loss of CDHR2 from intestinal and kidney epithelia in a villin-Cre driven knockout (KO) mouse, causes shortening and loss of brush border microvilli, a consequential decrease in the apical enrichment of key solute transporters, and reduced animal growth rate [14].

How new microvilli assemble and incorporate into a highly ordered brush border during differentiation remains unclear. Ultrastructural studies of native tissue and time-lapse imaging of epithelial cell culture models indicate that microvilli do not grow synchronously, but instead appear stochastically on the apical surface throughout differentiation [8, 11, 20]. One critical factor that promotes microvillar growth is the barbed end binder, epidermal growth factor receptor pathway substrate 8 (EPS8) [20, 21]. Previous studies in multiple epithelial and non-epithelial systems have established that EPS8 is a highly specific marker of the distal ends of all forms of actin bundle supported protrusions [22–25]. Loss of this factor leads to shorter protrusions and increased length variability [26, 27]. Strikingly, on the apical surface of differentiating epithelial cells, EPS8 arrives in diffraction-limited puncta at the membrane minutes before the subsequent growth of a core actin bundle and assembly of a microvillus at these sites [20]. Even once a core bundle begins to elongate, EPS8 puncta remain persistently associated with the distal end of the nascent structure. Following their initial growth, nascent microvilli are highly motile and translocate across the apical surface via a mechanism powered by treadmilling of the underlying core actin bundle [28], an activity that is also regulated by EPS8 [28]. Remarkably, if the distal tip of a newly formed microvillus loses its EPS8 punctum, that structure rapidly collapses, suggesting that EPS8 serves as a microvillus survival factor [20]. These data also point to a previously unrecognized dynamic microvillus lifecycle, consisting of distinct phases of structural stability and instability. For microvilli to eventually accumulate in large numbers on the apical surface, this cycle must ultimately tilt in favor of stability. However, how dynamic, nascent microvilli are stabilized on the apical surface so that they eventually accumulate long-term remains unknown.

Here we report our discovery of an adhesion-based mechanism that epithelial cells use to stabilize and in turn, drive the accumulation of microvilli during differentiation. Because microvillar growth takes place as differentiating enterocytes move through the crypt-villus transition [8], we reasoned that we could gain insight on mechanisms of microvilli accumulation by careful inspection of apical morphology in this region. Using this approach, we discovered that crypt microvilli initially accumulate at cell margins, implying the existence of a mechanism for anchoring nascent protrusions at these sites. We observed similar marginal accumulation of microvilli on the surface of differentiating intestinal and kidney epithelial cell lines. In all models examined, microvilli extending from one cell span intercellular space to make physical contact with microvilli on a neighboring cell. Using super-resolution microscopy, mechanistic studies in epithelial cell culture models, and live imaging, we determined that these points of physical contact represent “*transjunctional* IMACs” containing both CDHR2 and CDHR5, which are highly stable complexes that capture nascent microvilli and constrain their motion. Consistent with this point, long-term live imaging revealed that microvilli accumulation at cell margins outpaces accumulation in medial regions of the surface early in differentiation. Thus, microvilli extending from neighboring epithelial cells participate in a novel form of epithelial cell-cell contact that promotes apical surface maturation. The adhesion-based capture mechanism reported is likely to inform our understanding of apical morphogenesis in other epithelial cell types that build surface specializations.

## RESULTS

### Differentiating epithelial cells exhibit a marginal enrichment of microvilli

To begin to understand how microvilli are stabilized and accumulate in large numbers during differentiation, we first examined the distribution of nascent protrusions early in the maturation process. To this end, we used scanning electron microscopy (SEM) to survey the apical surface of the crypt cells in fractured samples of mouse small intestine. Within the crypt, where immature enterocytes are actively assembling a brush border (**Fig. 1A, zoom**), we noted a striking enrichment of microvilli at cell margins (**Fig. 1B, zoom 1 and 2 blue outlines**). In contrast, medial regions of the apical surface presented only a few, sparse microvilli (**Fig. 1B)**. Thus, *in vivo*, microvilli appear to accumulate at the edges of cells during the early stages of brush border assembly.

**Figure 1:**
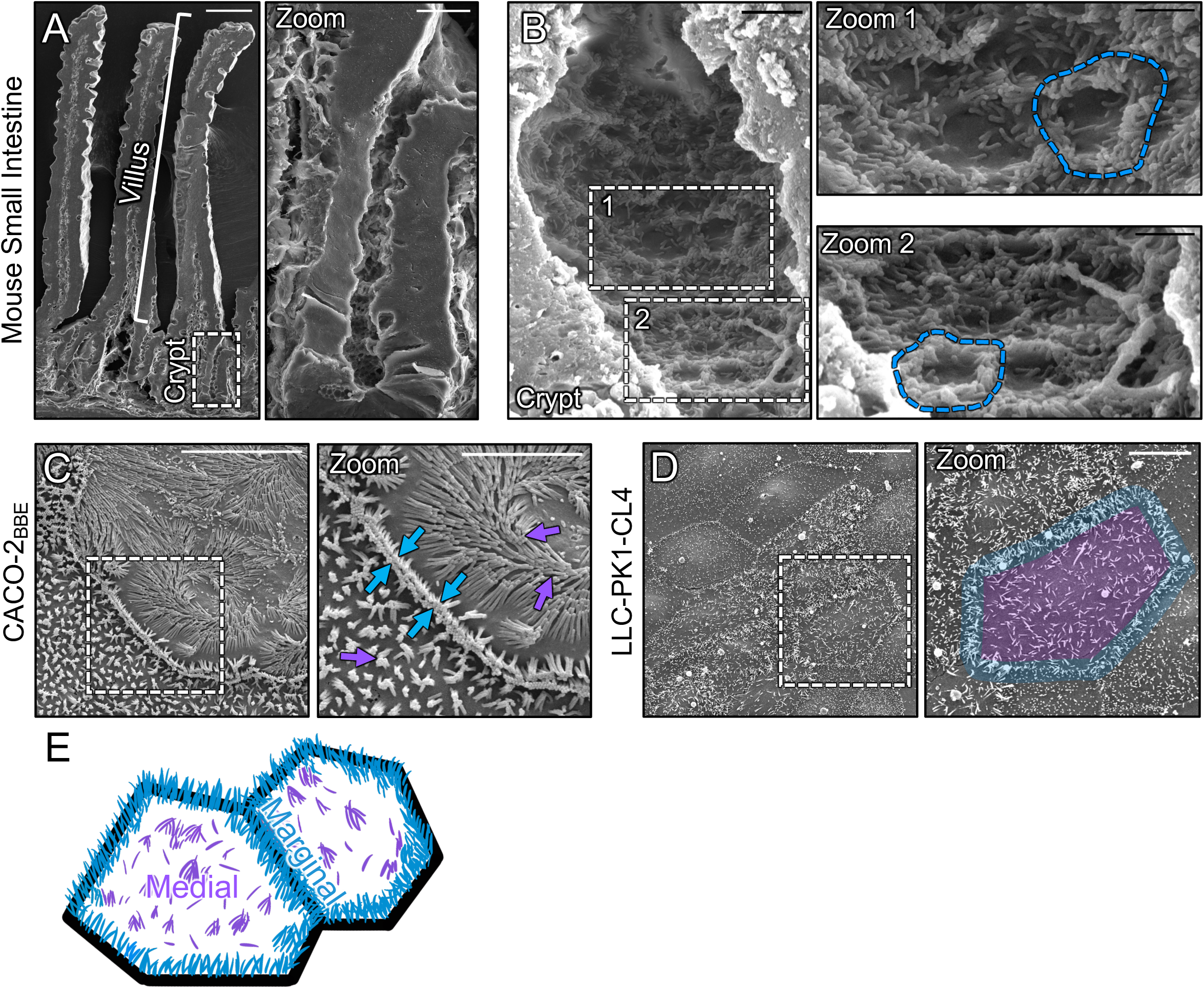
Microvilli of differentiating transporting epithelial cells concentrate at cell margins. (**A**) Scanning electron micrograph (SEM) of native mouse small intestine crypt-villus axis. (**A, zoom**) Zoom of the dashed box in A showing the crypt and transit amplifying zone. **(B**) High-magnification view of the crypt base with (**B, zooms 1 and 2**) showing an enrichment of microvilli at the margins of crypt cells (dashed blue outline). (**C**) SEM of polarized CACO-2_BBE_ cells. Dashed box represents zoom area. Arrows denote medial (purple) and marginal microvilli (blue). (**D**) SEM of sub-confluent porcine kidney proximal tubule LLC-PK1-CL4 (CL4) cells. Dashed box represents zoom area. Pseudo coloring represents medial area (purple) and marginal microvillar area (blue). (**E**) Schematic of the two distinct organizations of microvilli found on differentiating transporting epithelial cells, medial (purple) and marginal (blue). Scale bars: 50 µm (A), 10 µm (A, zoom), 2 µm (B), 1 µm (B, zooms), 10 µm (C), 5 µm (C, zoom), 20 µm (D), 10 µm (D, zoom).

To determine if the marginal accumulation of microvilli that we observed on the surface of differentiating crypt cells *in vivo* could be recapitulated *in vitro*, we first turned to the CACO-2_BBE_ line. CACO-2_BBE_ cells are a human intestinal epithelial cell culture model that builds a well-organized brush border over the course of several weeks at post-confluency [29]. SEM imaging of CACO-2_BBE_ cells early in the differentiation time-course revealed a concentration of microvilli at cell margins similar to that observed in native crypts (**Fig 1C**). Moreover, protrusions in these regions appeared to span cell junctions and make physical contact with structures on neighboring cells (**Fig. 1C, zoom blue arrows)**. As an additional point of comparison, we also examined SEM images of sub-confluent porcine kidney proximal tubule LLC-PK1 clone 4 (CL4) cells [30], which also exhibited a marginal accumulation of microvilli (**Fig. 1D, zoom blue outline**), even at the earliest stages of cell surface organization (i.e. subconfluence). Based on these *in vivo* and *in vitro* observations, the differentiating apical surface is characterized by two distinct populations of microvilli, marginal vs. medial **(Fig. 1E)**, with the marginal region demonstrating higher protrusion packing density at these early time points.

### Microvilli adopt a vertical orientation upon arriving at cell margins

In the ultrastructural images alluded to above, we noted that marginal microvilli appeared more vertically oriented relative to microvilli extending from medial parts of the cell surface. Here we use ‘vertical’ to describe an orientation that is parallel to the long (apicobasal) axis of the cell and perpendicular to the plane of the apical surface. To confirm this observation under hydrated conditions, we performed volume imaging of live sub-confluent CL4 cells expressing mCherry-Espin (ESPN), which serves as a highly specific marker of microvillar core actin bundles [20, 28, 31, 32] (**Fig. 2A**). Lateral viewing of reconstructed volumes enabled us to visualize individual microvilli and obtain measurements of their orientation relative to the plane of the apical surface. This analysis revealed that the marginal and medial microvilli demonstrate significant differences in their angle of protrusion, with marginal microvilli exhibiting a more vertical orientation (46.5° ± 19.3° vs. 77.3° ± 12.4°, **Fig. 2B**).

**Figure 2:**
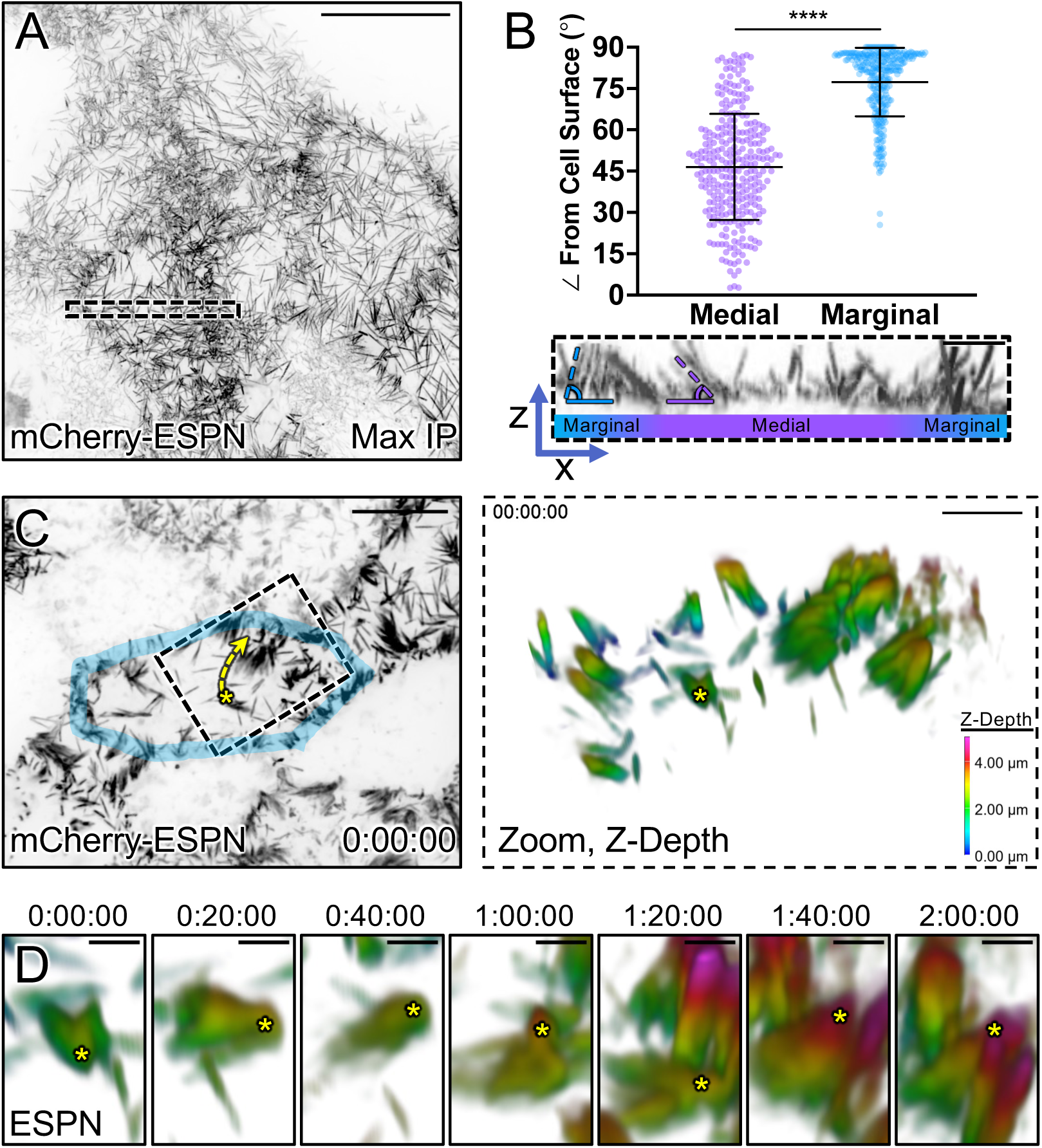
Microvilli adopt a vertical orientation upon reaching cell margins. (**A**) Maximum intensity projection (MaxIP) of live CL4 cells expressing mCherry-ESPN. (**B**) Orientation measurements of the angle (dashed outlines) of microvilli to the cell surface of medial microvilli (purple) compared to marginal microvilli (blue). Sample ROI of Z-projection under plot is taken from the dashed box in (A). (**C**) t = 0 MaxIP image of live mCherry-ESPN CL4 cells. One cell margin is highlighted in blue, while the dashed yellow arrow represents the trajectory of the microvilli cluster (asterisk) shown in (D). Right panel shows a 3D tilted volume of the dashed box in (C), coded in Z for cell depth (see Z-depth key on bottom right). (**D**) Montage over 2 hours following the cluster marked with the yellow asterisk in (C). Asterisk marks the distal ends of microvilli that transition to a vertical orientation upon reaching the marginal cell area, as shown by a change in Z-depth coding. Each point on the graph represents one angle taken from 17 cells; total of *n = 295* medial and *n = 309* marginal angles. Error bars represent mean ± SD. **** p ≤ 0.0001 Welch’s unpaired t-test. Mean medial angle is 46.5° ± 19.3° and mean marginal angle is 77.3° ± 12.4°. Scale bars: 20 µm (A), 1 µm (B), 10 µm (C, left), 3 µm (C, right), 1 µm (D).

Previous studies established that nascent microvilli are highly dynamic, growing, collapsing, and adopting a range of angles while undergoing active movement across the medial cell surface [20, 28]. With this in mind, we next sought to determine if microvilli grow in a vertical orientation at marginal sites or instead, grow medially and then adopt a vertical orientation upon arriving at the cell edge. To this end, we performed multi-hour time-lapse volume imaging to record microvillar motion and orientation in 3D. To help us interpret these complex datasets, we depth-coded volumes with a multi-color look-up table (LUT) so that image planes located further from the apical surface were rendered with warmer colors. While the dense accumulation of microvilli at cell margins impaired our ability to resolve individual growth events at these sites, we did observe individual protrusions and small adherent clusters of microvilli migrating while maintaining a small angle relative to the medial apical surface, as previously described [20, 28]. Following these microvilli over time (> 2 hrs) revealed that upon reaching the cell margin, they become more vertically orientated as indicated by the distal tips acquiring a warmer color coding (**Figs. 2C,D and Video S1**). Although these data do not allow us to rule out the possibility that microvilli grow *de novo* in a vertical orientation in the marginal zone, they do indicate that medial microvilli can transition into the marginal zone and adopt a vertical orientation upon doing so.

### Marginal microvilli are less motile than medial microvilli

Vertically orientated microvilli are a defining feature of mature brush borders on the surface of villus enterocytes [3]. Based on this point, the vertically oriented microvilli found in the marginal zone may represent more mature, and potentially, more stable protrusions. To begin to test this concept, we expressed EGFP-EPS8 to specifically mark the distal tips of microvilli [20, 21] in CL4 cells also expressing mCherry-ESPN (**Fig. 3A, zooms**). We then performed live volume imaging with the goal of using the punctate and stoichiometric EPS8 signal (one punctum per microvillus) as a high-fidelity fiducial marker for tracking microvillar dynamics over time. Temporal color coding of the ESPN channel over the course of 25 minutes revealed that medial microvilli are highly dynamic and demonstrate extensive movement as previously reported [28] (**Fig. 3B, zoom 1**). In contrast, marginal microvilli appeared to dwell for long periods near the edge of the cell, as indicated by the white band of color (merged colors of time points 0-25 min) in the projection (**Fig. 3B, zoom 2**). Next, we tracked individual EGFP-EPS8 puncta and generated rose plots of the resulting trajectories to examine the extent of motion demonstrated by individual microvilli. This analysis revealed that medial microvilli produce long trajectories consistent with directed motion, sampling an area of up to 6 µm^2^ during the time-lapse period (**Figs. 3C,D**). By comparison, the trajectories of marginal microvilli were highly confined, with individual protrusions traveling less than 2 µm^2^ over the same time course (**Figs. 3F,G**). Mean squared displacement analysis of trajectory data [28] confirmed that marginal microvilli are ∼10-fold more constrained in their movement (**Fig. 3H**) relative to medial protrusions (**Fig. 3E**). Together, these data suggest the existence of a mechanism for restricting the motion of microvilli at cell margins.

**Figure 3:**
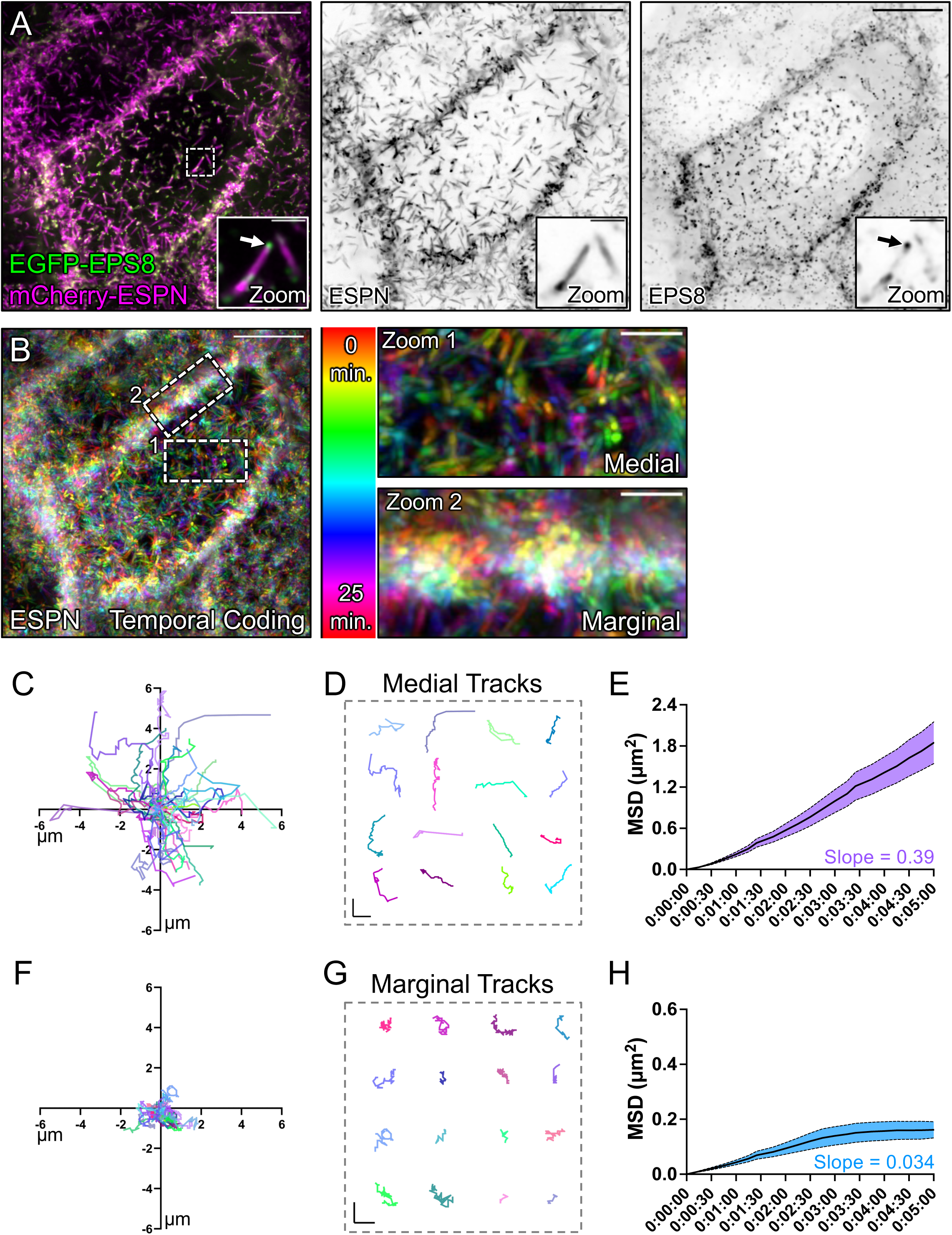
Tip tracking analysis reveals that marginal microvilli are constrained in their movement. (**A, left panel**) Live CL4 cells co-expressing EGFP-EPS8 and mCherry-ESPN. Dashed box represents zoom area with arrow marking EPS8 at the tip of a single microvillus. (**A, right panels**) Single inverted channel MaxIP images showing mCherry-ESPN and EGFP-EPS8 alone. (**B**) Temporal color-coding over 25 minutes (see vertical color key). (**B, zooms**) of (**1**) medial and (**2**) marginal ROIs taken from the dashed boxes in (B). (**C**) Rose plot of *n = 53* XY tracks (µm units) of medial microvilli over 25 minutes. (**D**) Representative medial microvilli tracks taken from (C). (**E**) Mean square displacement of 50 medial microvilli imaged for 5 minutes over 15 second intervals. Slope = 0.39. (**F**) Rose plot of *n = 28* XY tracks of marginal microvilli taken from 3 independent live cell imaging experiments over 25 minutes. (**G**) Representative marginal microvilli tracks taken from (F). (**H**) Mean square displacement analysis of *n = 88* marginal microvilli imaged for 5 minutes over 15 second intervals. Slope = 0.034. Scale bars: 10 µm (A), 1.5 µm (A, zooms), 10 µm (B), 2.5 µm (B, zooms), 1 µm (D, G).

### Microvilli from neighboring cells are linked by transjunctional adhesion complexes containing CDHR2 and/or CDHR5

Our ultrastructural data suggested that marginal accumulations of microvilli might include protrusions from both cells of a neighboring pair (**Fig. 1C zoom**). This led us to consider the possibility that microvilli extending from one cell may span the junctional space and physically contact microvilli from an adjacent cell; such interactions might in turn explain the upright orientation, reduced motility, and eventual accumulation of microvilli at these sites. One potential mechanism for mediating such interactions involves the intermicrovillar adhesion complex (IMAC), which includes the protocadherins CDHR2 and CDHR5 as core components [11]. Previous studies established that CDHR2 and CDHR5 target to the distal tips of microvilli and interact with each other to form a Ca^2+^-dependent heterophilic extracellular adhesion complex that spans the ∼50 nm between adjacent protrusions [8–10]. The resulting link promotes the tight packing of neighboring microvilli and contributes to minimizing length variability throughout the larger structure of the brush border [16]. Notably, these previous studies on IMAC function focused solely on *medial* microvilli, so the possibility that this complex might also link microvilli from neighboring cells remains unexplored.

To test this idea, we used an immunostaining approach and super-resolution structured illumination microscopy (SIM) to examine the localization of CDHR2, CDHR5, and F-actin relative to ZO-1, a critical component of tight junctions [33]. For these studies, we first examined native small intestinal tissues isolated from a new mouse model expressing CDHR2 tagged with EGFP at the endogenous locus. SIM images revealed that both IMAC protocadherins are highly enriched at the tips of medial microvilli as previously reported (**Fig. 4A**) [11]. We also noted signal from CDHR2 and CDHR5 at the tips of microvilli at the margins of cells, with the adhesion protein signal spanning ZO-1 marked junctions (**Fig. 4B, top panel**). When viewing projected SIM volumes *en face*, we were unable to discern the position of the tight junctions based solely on the phalloidin, CDHR2, or CDHR5 signals, suggesting that the IMAC components form a continuous network that spans beyond the surface of a single cell (**Fig. 4B, bottom panel**). Similarly, on the surface of mature enterocytes viewed with electron microscopy, microvilli appear to associate at neighboring cell interfaces (**Fig. S1A-B**). However, the crowded nature of microvilli in these mature brush borders confounded our attempts to isolate and visualize interactions between the tips of individual protrusions at the margins of neighboring cells.

**Figure 4:**
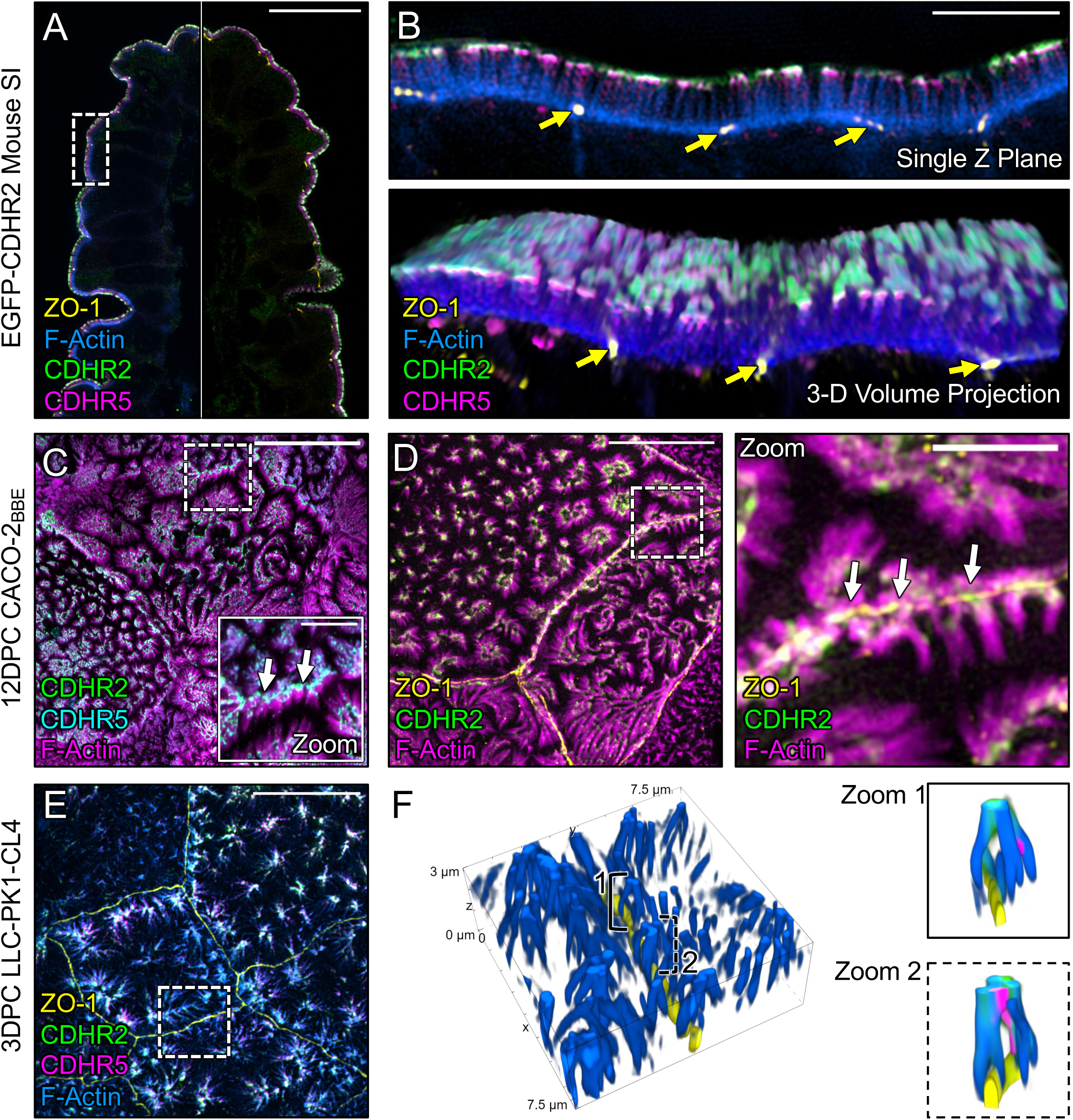
Marginal microvilli are linked via transjunctional CDHR2/CDHR5 adhesion complexes across neighboring cell junctions. (**A**) Single Z-plane confocal image of CDHR2-EGFP mouse small intestine stained for ZO-1 (yellow), EGFP (green), CDHR5 (magenta), and F-actin (blue). (**B, top**) Single plane SIM image of the stained villus section; approximated area marked by the dashed box in (A). (**B, bottom**) 3D volume projection of the top panel. Yellow arrows in both images mark ZO-1 labeled tight junctions. (**C**) MaxIP laser-scanning confocal image of 12 DPC CACO-2_BBE_ cells stained for CDHR2 (green), CDHR5 (cyan), and F-actin (magenta). Dashed box represents zoom area. White arrows point to tip-localized CDHR2/CDHR5 adhesion complexes at cell margins. (**D**) MaxIP SIM image of 12DPC CACO-2_BBE_ cells stained for ZO-1 (yellow), CDHR2 (green), and F-actin (magenta). Dashed box represents zoom area. White arrows point to CDHR2/CDHR5 marked complexes at the junction of neighboring cells. (**E**) MaxIP SIM image of 3 DPC CL4 cells stained for ZO-1 (yellow), CDHR2 (green), CDHR5 (magenta), and F-actin (blue). (**F**) 3D tilted volume projection of the dashed box outlined in (E). Brackets highlight instances of marginal microvilli on adjacent cells linked via CDHR2/CDHR5 transjunctional adhesion complexes (**zoom 1 and 2**, respectively). Scale bars: 20 µm (A), 5 µm (B), 2.5 µm (C), 5 µm (C, zoom), 10 µm (D), 2.5 µm (D, zoom), 10 µm (E).

To work around the limitation imposed by microvillar crowding in native tissue, we used SIM to examine the apical surface of cultured CACO-2_BBE_ cells at 12 days post-confluence (DPC), a time point *before* brush border assembly is complete, when microvillar packing density is comparatively lower. Careful examination of phalloidin-stained CACO-2_BBE_ monolayers revealed a striking enrichment and alignment of microvilli at the margins of cells (**Fig. 4C,D**), consistent with the SEM images described above (**Fig. 1B**). Immunofluorescence staining of these 12 DPC cultures revealed that marginally aligned microvilli do in fact span the cell junction marked by ZO-1 and exhibit enrichment of both protocadherins at their distal tips (**Fig. 4C,D, white arrows**). We observed similar structures and staining on the surface of CL4 monolayers at 3 DPC, a stage in differentiation when microvilli are still sparse but begin to form clusters and demonstrate marginal alignment (**Fig. 4E**). In this case, super-resolution lateral views clearly showed that individual microvilli from neighboring cells span the ZO-1-labeled tight junction and make contact via their distal tips, which are marked by both CDHR2 and CDHR5 (**Fig. 4F zooms**). In combination, these results indicate that marginal microvilli on neighboring cells are physically linked via *transjunctional* IMACs that contain CDHR2 and CDHR5.

### Heterophilic adhesion between CDHR2 and CDHR5 promotes robust association between microvilli from neighboring cells

Although IMAC protocadherin adhesion properties differ across species [16], previous biochemical studies established that in humans, heterophilic complexes of CDHR2 and CDHR5 exhibit strong adhesion, CDHR2 demonstrates weak homophilic adhesion, and CDHR5 demonstrates no homophilic adhesion [11]. To further study the nature of transjunctional IMACs, we developed a cell mixing approach that enabled us to drive the formation of adhesion complexes consisting of different complements of CDHR2 and/or CDHR5 (**Fig. 5A**). For these experiments, we first transfected CL4 cells with either EGFP or mCherry-tagged constructs of *H. sapiens* CDHR2 and CDHR5. Stable selection and subsequent fluorescence-activated cell sorting (FACS) yielded robust populations of fluorescent protocadherin expressing cells (**Fig. S2A**). Strikingly, mixed monolayers composed of cells expressing CDHR2-EGFP or CDHR5-mCherry demonstrated robust alignment of protocadherin signals at mixed cell-cell contacts **(Fig. 5B)**. Linescan analysis also revealed that CDHR2 and CDHR5 intensities were well correlated (mean r = 0.70) along these interfaces (**Fig. 5C,D,K**). These data are consistent with the formation of heterophilic adhesion complexes between microvilli of neighboring cells. In mixed monolayers composed of cells expressing CDHR2-EGFP or CDHR2-mCherry (**Fig. 5E**), mixed cell-cell contacts lacked the strong alignment of signals that we observed in the heterophilic case, and protocadherin intensities were poorly correlated (mean r = 0.07 (**Fig. 5F,G,K**). Mixed monolayers composed of cells expressing CDHR5-EGFP or CDHR5-mCherry also demonstrated a lack of signal alignment and poor intensity correlation along cellular junctions (mean r = -0.19) (**Fig. 5H-K**). High-resolution imaging of the interfaces formed under each of these three conditions revealed that only heterophilic mixtures of cells expressing CDHR2-EGFP or CDHR5-mCherry aligned their microvilli at cell-cell contacts (**Fig. S2B-D, white arrows**). Based on these data, we conclude that heterophilic transjunctional IMACs containing CDHR2 and CDHR5 can drive robust interactions between microvilli extending from neighboring cell margins.

**Figure 5:**
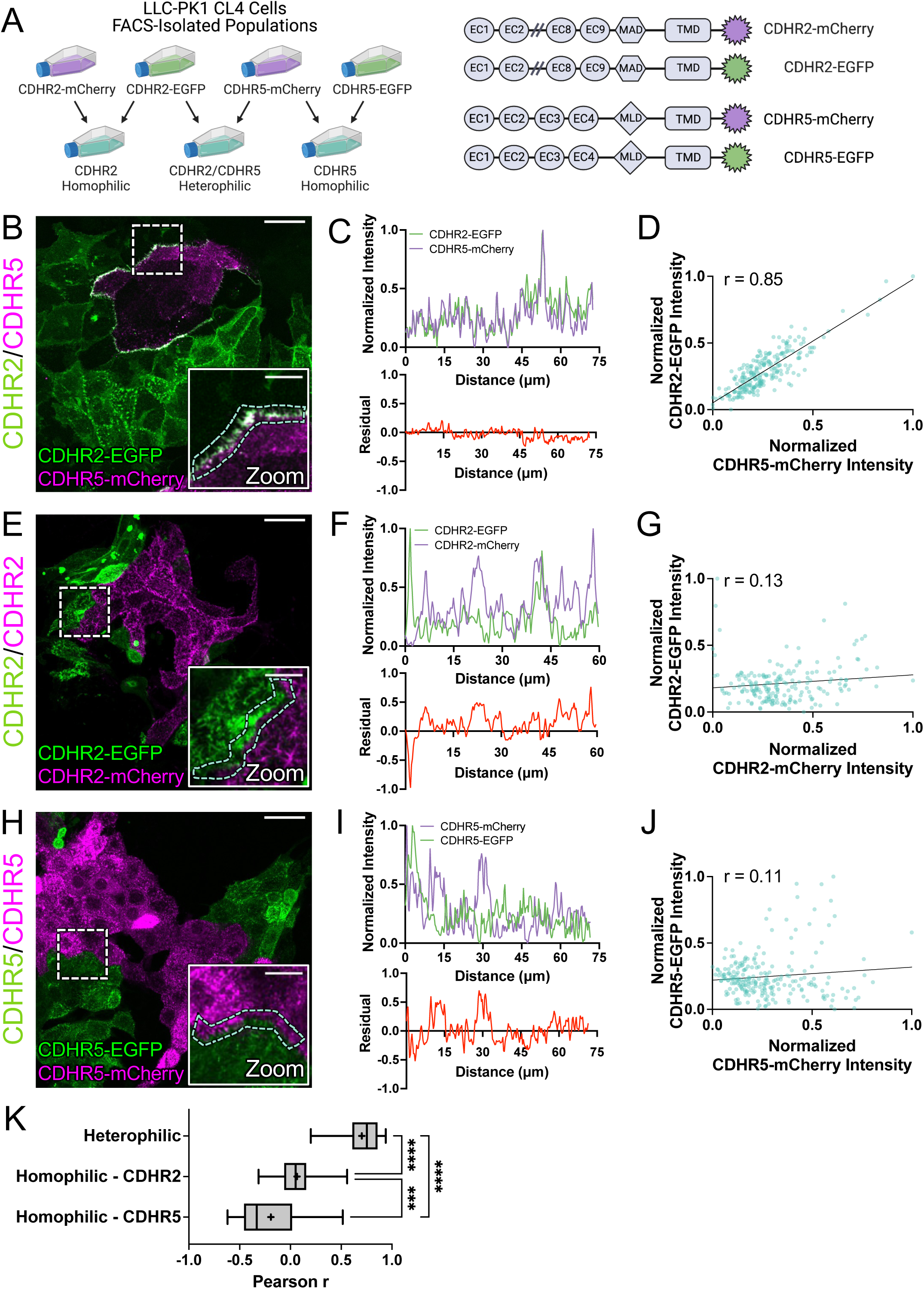
Cell mixing experiments reveal robust heterophilic adhesion complexes between marginal microvilli. (**A, left**) Schematic depicting cell mixing method for the C-terminally tagged cadherin overexpression constructs (**A, right**). (**B**) MaxIP laser scanning confocal image of mixed heterophilic CDHR2-EGFP and CDHR5-mCherry CL4 cell populations. Dashed box represents zoom area and cyan dashed outline represents sample linescan. (**C**, **top**) Normalized fluorescence intensity (AU) plot taken from a representative linescan along the mixed cell interface. (**C, bottom**) Plotted difference (residual) of mCherry signal from EGFP signal from the top linescan plot. (**D**) Pearson’s r correlation plot from the linescan in (C); r = 0.85. (**E**) MaxIP of mixed homophilic CDHR2-EGFP and CDHR2-mCherry CL4 cells. (**F-G**) Representative linescan and respective Pearson’s r correlation; r = 0.13. (**H**) MaxIP of mixed homophilic CDHR5-EGFP and CDHR5-mCherry CL4 cells. (**I-J**) Representative linescan and respective Pearson’s r correlation; r = 0.11. (**K**) Combined Pearson’s r values from *n = 30* individual linescans of each cell mixing scenario from 3 independent fixation and staining experiments (10 linescans per experiment). Mean Pearson’s r values are denoted by a “+” for each scenario where heterophilic r = 0.703, homophilic CDHR2 r = 0.065, and homophilic CDHR5 r = -0.193. Ordinary one-way ANOVA with multiple comparisons; **** p ≤ 0.0001 and *** p ≤ 0.001. Scale bars: 30 µm (B, E, H), 10 µm (zoom insets).

### Protocadherins in transjunctional IMACs exhibit limited turnover

Under normal conditions, epithelial cells express both CDHR2 and CDHR5, which target to the tips of all microvilli on the apical surface. Thus, heterophilic complexes are expected to form between the distal tips of microvilli in both the medial and marginal regions. However, the strong alignment of microvilli at cell-cell contacts in the heterophilic case outlined above led us to predict that transjunctional IMACs may be more stable relative to complexes that form medially. If true, this would offer a mechanistic explanation for the reduced motility of marginal microvilli, and in turn, the accumulation of microvilli at these sites. To determine if transjunctional IMACs are in fact longer lived than medial complexes, we performed fluorescence recovery after photobleaching (FRAP) analysis with CL4 monolayers formed using the cell mixing approach outlined above (**Fig. 5A**). Strikingly, photobleached ROIs positioned over junctional interfaces between heterophilic CDHR2-EGFP and CDHR5-mCherry expressing cells demonstrated extremely low signal recovery for both protocadherins (immobile fractions, 0.71 and 0.85, respectively; **Figs. 6A,B and Video S2**). In contrast, FRAP analysis of medially positioned ROIs on individual cells expressing both CDHR2-HALO and CDHR5-EGFP, revealed much lower immobile fractions for both protocadherins (0.47 and 0.56, respectively; **Figs. 6C,D and Video S3**). These results suggest that transjunctional IMACs formed between marginal microvilli are much longer lived relative to complexes formed between the tips of medial microvilli. We also examined recovery in photobleached ROIs positioned over junctional interfaces formed between homophilic CDHR2-EGFP and CDHR2-mCherry expressing cells (**Figs. 6E,F and Video S4**), as well as interfaces formed between homophilic CDHR5-EGFP and CDHR5-mCherry expressing cells (**Figs. 6G,H and Video S5**). Both homophilic scenarios exhibited higher levels of turnover and even lower immobile fractions. Together, these FRAP studies indicate that transjunctional IMACs composed of CDHR2 and CDHR5 are extremely stable, and these reduced turnover kinetics offer a explanation for the constrained motility and accumulation of microvilli observed at cell margins.

**Figure 6:**
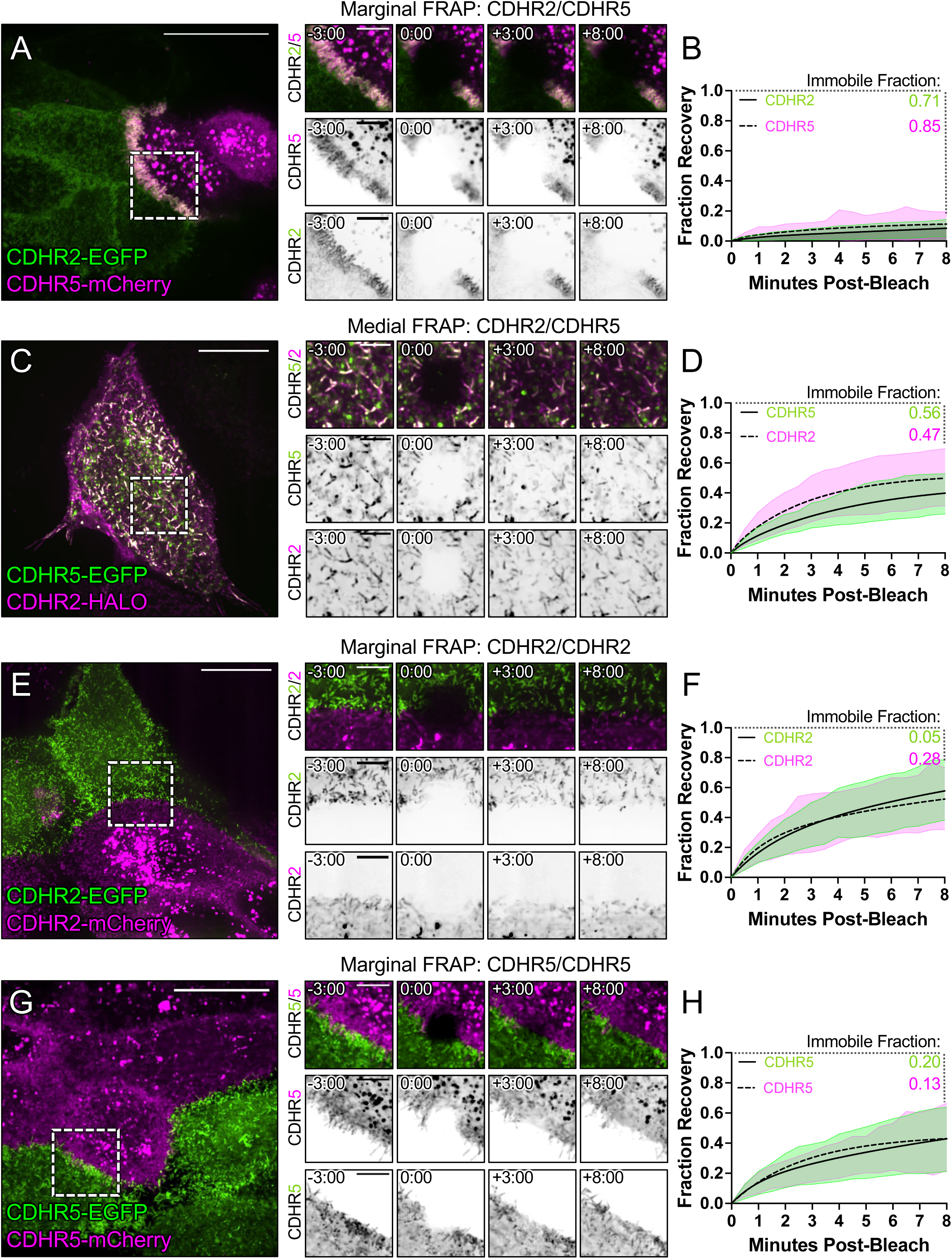
FRAP analysis suggests that heterophilic, transjunctional adhesion complexes are stable. Mixed CL4 cells forming (**A**) marginal heterophilic, (**C**) medial heterophilic, (**E**) marginal homophilic CDHR2, and (**G**) marginal homophilic CDHR5 adhesion complex interfaces. Dashed boxes outline the photobleached ROI shown in the recovery montages on right. (**B, D, F, H**) Fluorescence recovery is plotted over the course of 8 minutes with the immobile fractions as written for each protein channel. All plots represent 3 independent FRAP experiments of *n ≥ 20* ROIs from multiple cells. Scale bars: 20 µm (A, C, E, G), 5 µm (montages).

### Microvillar packing density at cell margins is higher than the medial zone during differentiation

Based on the stabilizing nature of transjunctional IMACs, we predicted that, during differentiation, cells might assembly the brush border by packing microvilli inward from cell margins. To test this idea, we performed extended time-lapse imaging of CL4 cells expressing mCherry-ESPN to stoichiometrically label microvillar core actin bundles (**Fig. 7A-C**)[32]. Comparing regional ESPN intensities on a per cell basis, we found that marginal ESPN intensity increased almost ∼2-fold more than medial signal during 24 hrs of differentiation (**Fig. 7D**). These timelapse results are consistent with the idea that microvilli accumulate first at cell margins and then pack inwards from the edges of the cell over time.

**Figure 7:**
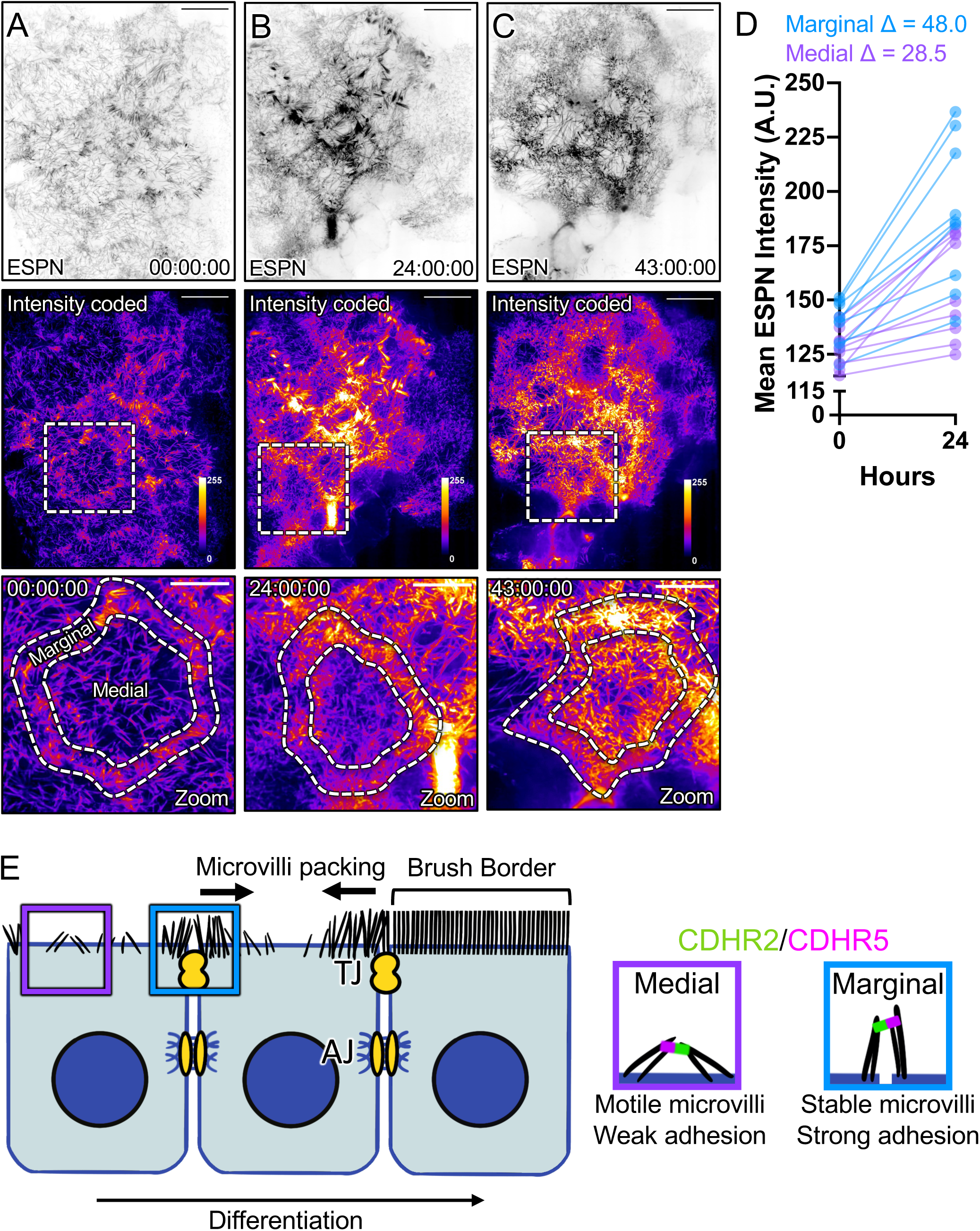
Long-term imaging reveals that microvilli first accumulate at cell margins over the course of differentiation. (**A-C**) MaxIP spinning disk confocal stills of live mCherry-ESPN expressing CL4 cells at t = 0 hours, 24 hours, and 43 hours from a 43-hour acquisition. (**A-C, intensity coded**) Fire LUT intensity profile of the mCherry-ESPN channel. Intensity scales from low (0; dark purple) to high (255; yellow/white) as denoted by LUT profile. Zooms at each time point are outlined by dashed boxes, with marginal and medial zones as marked. (**D**) Mean marginal and medial mCherry-ESPN intensity of *n = 10* cells from the time points shown in (A-C). Change in mean ESPN intensity (AU) in the first 24 hours is denoted on the graph. The cut axis accounts for background mCherry signal. Scale bars: 20 µm (A-C), 10 µm (zooms). (**E**) An adhesion-based model for the marginal stabilization of microvilli during brush border assembly. Microvilli on nascent transporting epithelial cells organize into two distinct populations: medial and marginal. Medial microvilli are highly motile while marginal microvilli are stable and stand at an orientation more vertical to the apical surface. Transjunctional CDHR2/CDHR5 heterophilic adhesion complexes span cell-cell junctions and link marginal microvilli of neighboring cells. These complexes are long-lived, and lead to the accumulation of microvilli at the edges of cells. A predicted outside-in packing mechanism occurs during differentiation as a result of transjunctional adhesion complexes first stabilizing microvilli at cell margins.

## DISCUSSION

Previous live imaging studies of epithelial cells at times points early in differentiation established that actively growing and newly formed microvilli are highly motile and unstable, undergoing rapid cycles of growth and collapse [20, 28]. Those discoveries immediately led us to question how dynamic, nascent microvilli are stabilized long-term on the apical surface to enable their timely accumulation in large numbers (i.e., thousands) by the end of differentiation. We first sought to approach this question by examining the surface of the undifferentiated epithelial cells that line the interior of the intestinal crypt, where microvillar growth activity is high. Because the apical surface of cells in this region is not yet fully packed with protrusions, we were hoping to identify patterns in the distribution of nascent microvilli that might offer insight on underlying mechanisms of stability.

Peering into the crypt is technically challenging given the tight confines of this invaginated compartment. Indeed, almost all previous ultrastructural studies of this region have been limited to conventional transmission EM of ultrathin sections [34, 35], which are difficult to interpret in the absence of 3D context. We worked around this obstacle using a combination of tissue fracturing and scanning EM, which allowed us to visualize the apical surface of immature intestinal epithelial cells within the crypt. Inspection of these images revealed that microvilli preferentially accumulate near the cell periphery at this point in differentiation. Moreover, cell culture models from the intestine (CACO-2_BBE_) and kidney (CL4) also demonstrated robust marginal accumulation of microvilli early in their maturation time course, suggesting that such patterning is not a function of the unique cellular packing geometry found in the crypt, nor is it tissue specific.

Accumulation of microvilli at cell edges suggests that the marginal zone represents (i) a site of robust growth, (ii) a site of stabilization for nascent microvilli, or (iii) some combination of the two. Given the actin-rich junctional belt that surrounds the cell at the level of the terminal web [36], it seems reasonable to expect that microvilli may grow more readily in this location. Although previous live imaging studies of CL4 cells characterized the properties of individual microvillar growth events [20], those observations were limited to the medial regions of the cell where protrusion density is typically low; visualization of growth events in the marginal zone was confounded by the crowding of pre-existing microvilli in this region. While our data do not allow us to rule out the possibility that growth preferentially occurs at cell margins relative to medial regions, we were able to capture clear examples of clustered microvilli moving at a low angle relative to the cell surface, toward the edge of the cell and incorporating into the marginal population. Interestingly, these protrusions adopt the more vertical orientation of marginal microvilli upon reaching the cell edge. Because such upright orientation is a defining feature of microvilli in mature brush borders, the marginal population likely represents stabilized protrusions that persist into later stages of differentiation. Although we currently lack a method for tracking and measuring the lifetimes of individual microvilli over the course of days, our short-term tracking measurements using the tip marker, EPS8, confirm that marginal microvilli are less motile relative to medial microvilli. Indeed, using mean square displacement analysis as previously described [28], we found that marginal microvilli sample ∼10-fold less surface area per unit time relative to medial microvilli, which is consistent with a physical capture mechanism near the cell edge.

Earlier work established that medial microvilli on the surface of mature villus enterocytes employ the protocadherins CDHR2 and CDHR5 to form intermicrovillar adhesion complexes (IMACs) that link the distal tips of neighboring microvilli [11, 15]. Here we sought to test the possibility that IMACs form across cell junctions, between the protrusions that extend from neighboring cells. If so, this would offer a mechanistic explanation for the upright orientation and constrained motility that microvilli demonstrate at these sites, and potentially the long-term stabilization that enables microvillar accumulation on the apical surface in large numbers. Previous work in CACO-2_BBE_ cells, native mouse intestinal tissue, and X-ray crystallography all indicate that the interacting ectodomains of CDHR2 and CDHR5 are structurally capable of spanning gaps up to 63 nm wide [11, 16], suggesting that they could easily reach across the ∼15 nm tight junction between neighboring cells [37]. Indeed, in the current study, super-resolution imaging revealed that CDHR2 and CDHR5 span the intercellular space to form *transjunctional IMACs* that physically link marginal microvilli that extend from neighboring cells.

Does the formation of transjunctional IMACs explain the accumulation of microvilli at cell edges early in differentiation? If transjunctional IMACs are more stable and exhibit longer lifetimes relative to IMACs that form medially, this would certainly offer a mechanistic underpinning for the increase in microvilli density at these sites. To test this hypothesis, we employed a cell mixing approach that enabled us to induce the formation of both homophilic and heterophilic transjunctional IMACs, to enable further characterization of their properties. FRAP analysis of the turnover dynamics of these complexes revealed that heterophilic (CDHR2/CDHR5) transjunctional IMACs are much longer lived relative to homophilic (CDHR2/CDHR2) complexes. These results from live epithelial cells echo previous *in vitro* data suggesting that homophilic (CDHR2/CDHR2) complexes are much weaker than heterophilic (CDHR2/CDHR5) complexes [11]. Interestingly, when we examined the dynamics of heterophilic complexes formed between microvilli in the medial population, we noted that these also turned over at a much higher rate relative to transjunctional heterophilic (CDHR2/CDHR5) complexes. Thus, the significant differential stability of transjunctional vs. medial IMACs indicated by our FRAP studies offers a mechanistic rationale for the accumulation of microvilli at cell margins.

Why transjunctional IMACs are more stable than those formed elsewhere on the apical surface remains unclear, but possible explanations might be found in previous biophysical studies on the properties of non-covalent bonds. For example, when a tensile mechanical force is applied across a non-covalent bond formed between two proteins, the lifetime of that bond will be impacted in a way that depends on the structural nature of the bonding interface [38, 39]. “Slip bonds” react to loading with a dramatic shortening of lifetime, whereas “catch bonds” respond by increasing bond lifetime; “ideal bonds” exhibit minimal response to mechanical loading [38, 40, 41]. Direct physical measurements provide strong evidence for catch bond behavior in structurally diverse proteins, ranging from myosin motor domains to cell surface molecules such as integrins [42, 43]. Cadherins have been studied extensively in this regard and their bonding properties are complex. In the case of E-cadherin, adhesive interactions can exhibit slip *or* catch behavior depending on the conformation of the adhesive interface. In the canonical strand swapped conformation, E-cadherin exhibits slip bond behavior; while X-dimers of E-cadherin, which interact using a distinct extended structural interface demonstrate robust catch bond behavior [40]. In light of those findings, we speculate that IMACs also exhibit catch bond properties. By bridging across cell junctions, transjunctional IMACs may be subject to higher tensile loads and therefore exhibit increased adhesive lifetimes relative to IMACs that form elsewhere on the apical surface. Furthermore, as medial clusters of microvilli appear to move as a unit [28], their adhesive bonds may be under less tensile stress. Although rigorous testing of this concept must await future biophysical studies, it is important to note that, based on the recently solved structures of mouse and human CDHR2 and CDHR5 ectodomains [16], any catch bond behavior in the IMAC would emerge from a mechanism that is distinct from E-cadherin.

Given the adhesive capture of microvilli by stable transjunctional IMACs and our observations of marginal microvilli enrichment early on in cell surface differentiation, we speculated that the brush border assembly may favor packing from the margins of the apical surface, inwards (**Fig. 7E**). To test this idea, we turned to multi-day time-lapse imaging of CL4 cells expressing mCherry-ESPN as a marker for microvilli. As expected, we noted that the marginal ESPN intensity was initially higher than in the medial region. After 24 hours of differentiation, the marginal region also demonstrated ∼2-fold larger increase in signal relative to the medial zone, suggesting that microvillar packing density at the cell margin precedes packing of the interior apical surface. Moreover, intensity at the cell margin is consistently higher than medial signal, and both regions increase in intensity over the course of almost two days of observation. In future studies, it will be critical to confirm this observation in systems that more closely recapitulate the biology of the crypt-villus transition, such as intestinal organoids.

While previous work established that the IMAC is critical for maintaining brush border structure on mature enterocytes [14], the current study indicates a new role for this complex in apical surface maturation, by contributing to a novel form of cell-cell contact between microvilli of neighboring cells. In the intestinal tract and other transporting epithelial tissue, cell-cell contacts are essential for “barrier” function and the maintenance of physical compartmentalization. Interestingly, Crohn’s Disease patients exhibit a decrease in CDHR2 and CDHR5 mRNA expression [44] while also experiencing increased intestinal permeability [45]. Transjunctional adhesion complexes may also form an additional layer of protection against colonizing pathogens. Infection by related pathogens Enteropathogenic and Enterohemorrhagic *Escherichia coli* (EPEC and EHEC) is characterized by effacement of brush border microvilli and F-actin pedestal formation [46]. CDHR2 has been identified as one of the initial EHEC targets during infection, which results in a significant decrease in CDHR2 expression [47]. Past reports on EPEC infection also show both bacteria localization over cell junctions [46, 48]. In the future studies, it will be fascinating to explore new roles for transjunctional IMACs in maintaining epithelial barrier function in intestinal disease and infection.

## ACKNOWLEDGMENTS

The authors would like to thank all members of the M.J.T. laboratory for their constructive feedback. Special thanks goes to laboratory manager Suli Mao, for her work with mouse husbandry and genotyping. We acknowledge the Translational Pathology Shared Resource (NCI Cancer Center Support Grant 5P30CA68485-19) and also thank the Vanderbilt Genome Editing Resource (RRID: SCR_018826). Microscopy was performed in part through the VUMC Cell Imaging Shared Resource. Some SEM images were collected on a Zeiss Crossbeam funded by 1S10 OD028704. Flow Cytometry experiments were performed in the VUMC Flow Cytometry Shared Resource supported by the Vanderbilt Ingram Cancer Center (P30 CA68485) and the Vanderbilt Digestive Disease Research Center (DK058404). This work was supported by the NIH grants DK095811, DK125546, DK111949 (M.J.T.) and the Training Program in Developmental Biology 2T32HD007502-20 (C.S.C.)

## AUTHOR CONTRIBUTIONS

Conceptualization, C.S.C. and M.J.T.; Methodology, M.J.T., C.S.C., B.A.M. and E.S.K.; Validation, C.S.C.; Formal Analysis, C.S.C., J.B.S., and M.J.T.; Investigation, C.S.C., J.B.S., L.M.M., and B.A.M.; Writing, C.S.C. and M.J.T.; Visualization, C.S.C.; Supervision, M.J.T.; Project Administration, M.J.T.; Funding Acquisition, C.S.C. and M.J.T.; All authors contributed to revising the manuscript.

## INCLUSION AND DIVERSITY

We support inclusive, diverse, and equitable conduct of research.

## METHODS

### Key Resources Table

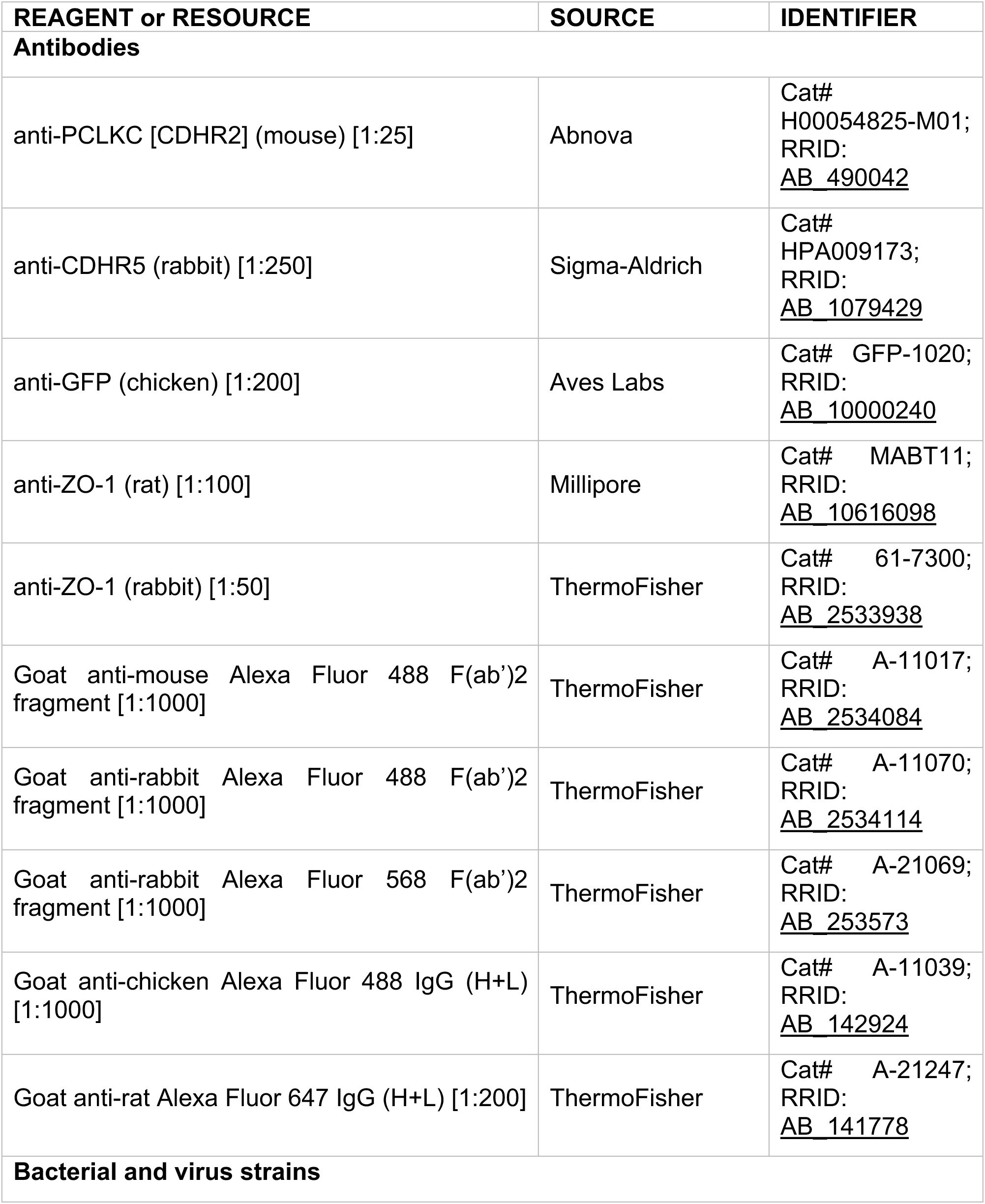
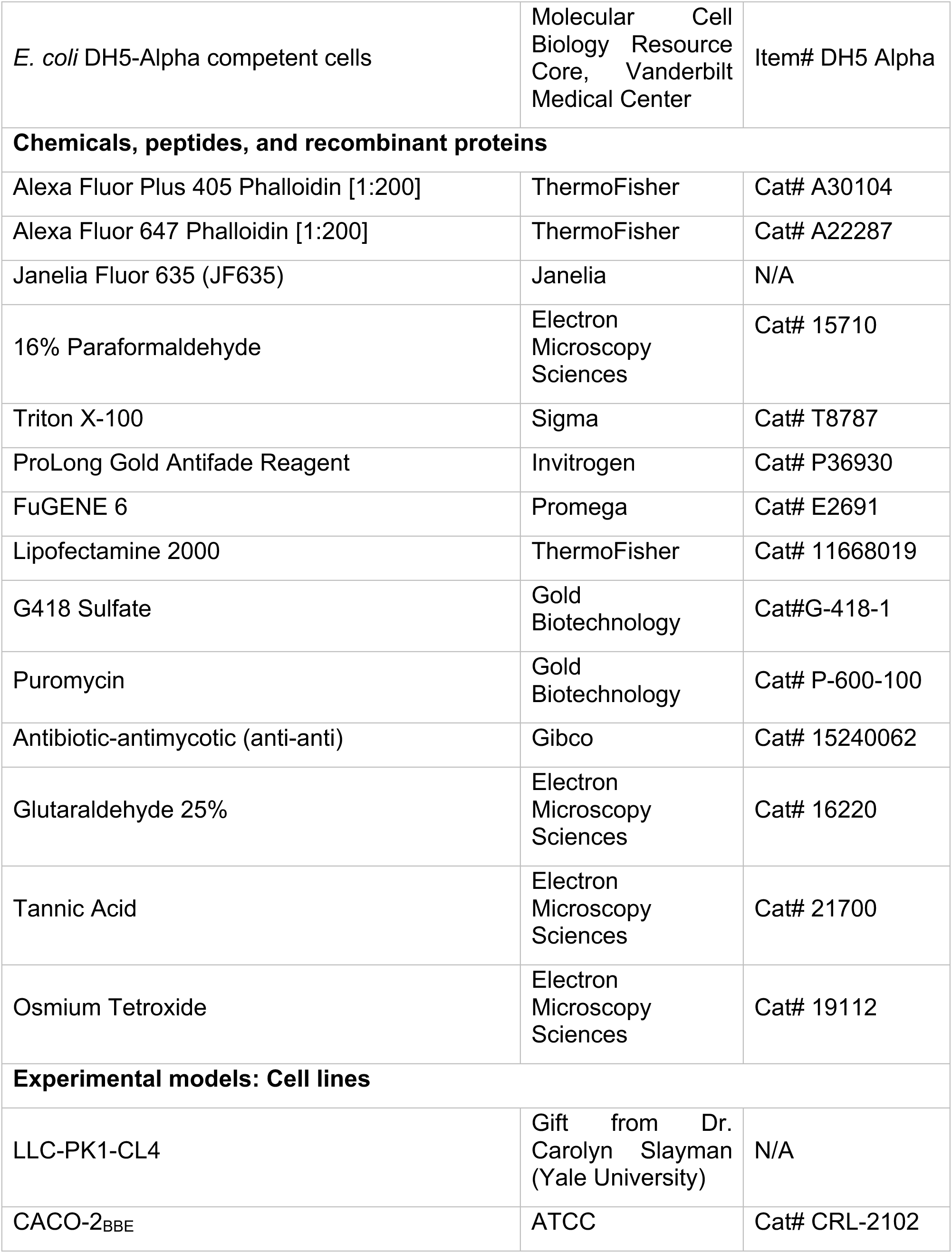
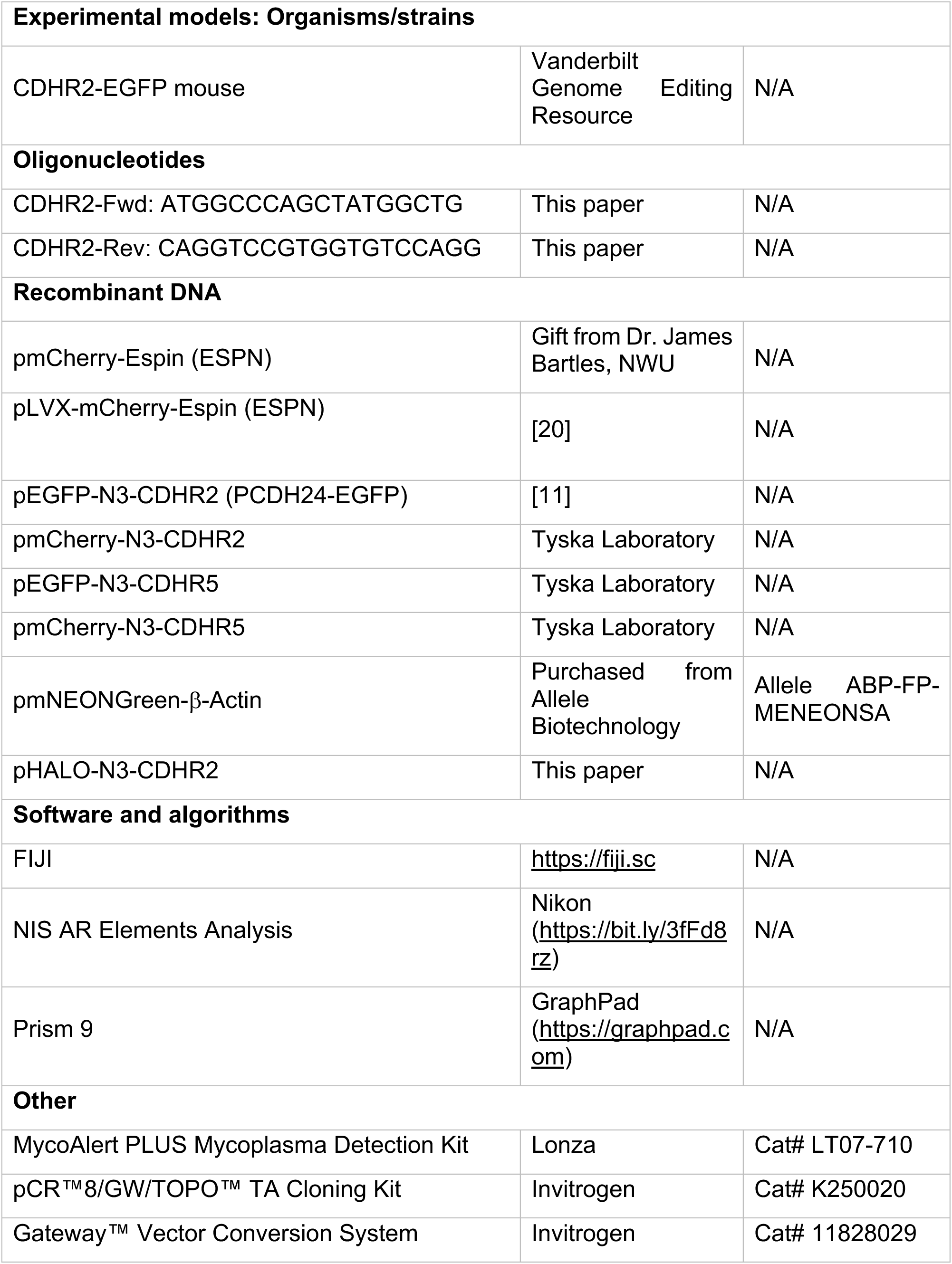
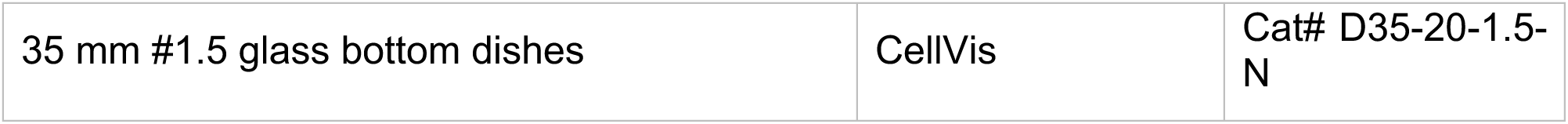

### Animal studies

Animal experiments were carried out in accordance with Vanderbilt University Medical Center Institutional Animal Care and Use Committee guidelines under IACUC Protocol ID#: M1600206-02.

### CDHR2-EGFP mouse

Created in collaboration with the Vanderbilt Genome Editing Resource. A C57Bl/6N strain containing a *CDHR2* C-terminal EGFP sequence insertion. [crRNA sequence: TGGACACCACAGATCTGTGA] Ribonucleoprotein complexes containing ctRNA and WT SpCas9 protein were targeted to the C-terminus of *CDHR2* were assembled and injected with a single stranded 944 nt DNA donor into 1-cell C57Bl/6N embryos. crRNA, tracrRNA, and WT SpCas9 protein was sourced from MilliporeSigma. The single stranded DNA was produced by Genewiz. Pups were screened for CDHR2-EGFP sequence insertions by PCR and validated by Sanger sequencing.

### Frozen Section Tissue Preparation

The proximal segment (duodenum to jejunum) of the mouse intestinal tube was excised and flushed with cold 1X phosphate-buffered saline (PBS). One end of the tube was clamped with a hemostat and the tube was filled with room temperature 2% paraformaldehyde (PFA) (Electron Microscopy Sciences) with a syringe and metal cannula. The other end of the tube was clamped with a hemostat and the tissue was laid in a petri dish containing excess 2% PFA and incubated for 15 minutes at room temperature. Hemostats were removed and the tissue was cut lengthwise into one flat piece. Tissue was then sub dissected into ∼2mm^2^ pieces and fixed for an additional 30 minutes in a vial of 2% PFA at room temperature. After fixation, the tissue was washed 3 times with PBS and then placed, villi-side down, into a vial of cold 30% sucrose/1% sodium azide. The tissue was placed at 4°C, overnight until sections sank to the bottom of the tube. The next day, sections were passed through 3 separate blocks of optimal cutting temperature (OCT) compound (Electron Microscopy Sciences) to wash off the sucrose solution, oriented with villi parallel to the lab bench in a fresh block of OCT, and snap frozen in dry ice-cooled acetone. Samples were cut into 10 µm thin sections using a Leica CM1950 cryostat and mounted on plasma-cleaned #1.5H precision coverslips (Thorlabs). Coverslips were stored at -20°C until staining.

### Frozen Section Immunofluorescence

Coverslips were thawed to room temperature and rinsed twice with 1X PBS to remove OCT. Sections were permeabilized with 0.2% Triton X-100 (diluted in PBS) for 10 minutes at room temperature. Sections were then rinsed once with PBS at room temperature and blocked in 10% bovine serum albumin (BSA) for 2 hours at 37°C in a humidified chamber. After rinsing with PBS, primary antibody (diluted in 1% BSA) was applied overnight at 4°C in a humidified chamber. The next day, sections were rinsed with 1X PBS 4 times and secondary antibody (diluted in 1% BSA) was applied for 2 hours at room temperature in a dark, humidified chamber. Sections were rinsed 4 times with 1X PBS and coverslips were mounted onto glass slides with ProLong Gold. The following antibodies and dilutions were used for EGFP-CDHR2 mouse frozen section staining: anti-GFP (chicken Aves #GFP-1020), 1:200; anti-CDHR5 (rabbit, Sigma #HPA009173), 1:250; anti-ZO-1 clone R40.76 (rat, EMD Millipore Sigma #MABT11), 1:100; Alexa Fluor goat anti-chicken 488 (Invitrogen #A-11039), 1:1000; Alexa Fluor F(ab’)2 fragment goat anti-rabbit 568 (Invitrogen #A21069), 1:1000; Alexa Fluor goat anti-rat 647 (Invitrogen #A21247), 1:200; and Alexa Fluor Plus 405 Phalloidin (Invitrogen #A30104), 1:200 for actin staining. The secondary antibodies, not including phalloidin, were spun down for 10 minutes at 4°C and 21 x g prior to using.

### Cell Culture

LLC-PK1-CL4 (porcine kidney proximal tubule) cells were grown in 1X high glucose DMEM containing 2mM L-glutamine (Corning #10-013-CV) supplemented with 1% L-glutamine (Corning # 25-005-CI) and 10% fetal bovine serum (FBS) (R&D Systems) while CACO-2_BBE_ (human colonic adenocarcinoma) cells were grown in the same medium but supplemented with 20% FBS. Cells were maintained in culture incubated at 37°C and 5% CO_2_. Cells were tested for mycoplasma using the MycoAlert PLUS Mycoplasma Detection Kit (Lonza #LT07-710).

### Cloning and Constructs

A C terminally tagged pHALO-N3-CDHR2 (CDHR2-HALO) construct was generated by taking full length CDHR2 via PCR from pEGFP-N3-PCDH24 (CDHR2-EGFP) with the primers CDHR2-Fwd: ATGGCCCAGCTATGGCTG and CDHR2-Rev: CAGGTCCGTGGTGTCCAGG. The product was then TOPO cloned into the pCR8/GW/TOPO vector (Invitrogen #K250020) and then placed into the pHALO-N3 backbone, adapted for Gateway cloning using the Gateway conversion kit (Invitrogen #11828029). All other overexpression constructs listed in this paper were previously created and/or reported as noted in the key resources table.

### Cell Line Generation

Cells expressing one plasmid were transfected with FuGENE 6 (Promega #E2691) at a FuGENE:DNA (μL:μg) ratio of 3:1 following the reagent protocol in a T25 cell culture flask. The next day, all cells were split up to a T75 flask with the addition of 1mg/mL G418 sulfate for antibiotic selection. Cells were maintained in culture under constant G418 selection to create a stably expressing cell line. Cells co-expressing two plasmids were transiently transfected with Lipofectamine 2000 (Invitrogen #11668019) according to the manufacturer’s protocol. The next day, cells were split to plasma-cleaned 35mm glass bottom dishes (CellVis #D35-20-1.5-N) for subsequent imaging. The EGFP-EPS8/mCherry-ESPN CL4 stable cell line was previously created [20] by transducing a G418-selected EGFP-EPS8 stable cell line with lentiviral mCherry-ESPN followed by 10 μg/mL puromycin selection. See citation for detailed protocol. The Halo-CDHR2/EGFP-CDHR5 co-expressing CL4 cells were a transient transfection, and not stably selected.

### Cell Immunofluorescence

Prior to fixation and staining, CL4 and CACO-2_BBE_ cells were grown to *n* days post-confluent (DPC) on acid-washed 22×22 mm #1.5H coverslips (Globe Scientific) in a 6-well plate to a time point with apical polarity representative of their native tissue, 3 DPC and 12 DPC, respectively. First, cells were rinsed in warm 1X PBS and fixed in 4% PFA for 15 minutes at 37°C. Cells were then washed three times, 5 minutes each, with 1X PBS and permeabilized with 0.1% Triton X-100 for 10 minutes at room temperature. 5% BSA was added for 1 hour at 37°C as blocking solution. After rinsing with 1X PBS, primary antibody (diluted in 1% BSA) was added for 1 hour at 37°C. Labeling with primary antibody was followed by washing 4 times, 5 minutes each, with 1X PBS. Secondary antibody (diluted in 1% BSA) was then applied for 1 hour at room temperature in the dark. After incubation in secondary antibody, cells were washed 4 times, 5 minutes each with 1X PBS and coverslips were mounted on glass slides with ProLong Gold. The following antibodies and dilutions were used for cell staining: anti-PCLKC (CDHR2) (mouse, Abnova #H00054825-M01), 1:25; anti-CDHR5 (Rabbit, Sigma #HPA009173), 1:250; anti-ZO-1 clone R40.76 in CL4 (rat, EMD Millipore Sigma #MABT11), 1:100; anti-ZO-1 in CACO-2_BBE_ (rabbit, Invitrogen #61-7300), 1:50; Alexa Fluor F(ab’)2 fragment goat anti-mouse 488 (Invitrogen #A11017) and goat anti-rabbit 568 (Invitrogen #A21069), 1:1000; Alexa Fluor goat anti-rat 647 (Invitrogen #A21247), 1:200; and Alexa Fluor Plus 405 Phalloidin (Invitrogen # A30104) or Alexa Fluor 647 Phalloidin (Invitrogen # A22287), 1:200 for actin staining. The secondary antibodies, not including phalloidin, were spun down for 10 minutes at 4°C and 21 x g prior to using.

### Fluorescence-activated Cell Sorting (FACS)

Cells were spun down into a pellet and resuspended in “pre-sort medium” containing Phenol free 1X DMEM (Gibco #21063-029) plus 5% FBS, and 1% L-glutamine. Cells were sorted by Vanderbilt University Medical Center’s Flow Cytometry Shared Resource on a 5-Laser FACS Aria III system with a 100 µm sized nozzle. All fluorescent positive cells (Fig. S3) were deposited into a single well of a 6-well plate containing “post-sort medium” of 1X DMEM (Corning #10-013-CV) with Phenol red, 10% FBS, 1% L-glutamine, and 10µL/mL anti-anti (Gibco #15240062). 24 hours post-sort, the media was changed to CL4 culture media (as detailed in cell culture methods) and 1 mg/mL G418 was added for maintaining stable plasmid overexpression. Sorted cell lines were maintained in this media and under antibiotic selection.

### Cell Mixing Experiments

Fluorescently sorted CL4 cell populations were grown independently and under G418 antibiotic selective pressure to ∼80% confluence, trypsinized, and resuspended in CL4 media to a density of ∼850,000 cells/mL. 250 µL of each cell population were seeded in plasma-cleaned glass-bottom dishes or onto acid washed coverslips at a density of ∼400,000 total cells at a mixing ratio of 1:1 (e.g. CDHR2-EGFP cells were mixed with CDHR5-mCherry cells). Immediately after seeding, cell populations were thoroughly mixed by pipette. Cells were grown to 3DPC for fixed cell staining or for 1 day for live cell imaging (FRAP).

### Fixed Sample Microscopy

<u>Laser scanning confocal microscopy</u> was performed on a Nikon A1 microscope equipped with 488 nm, 568 nm, and 647 nm LASERs. Mixed CL4 cell populations for linescan analysis were imaged using a Plan Apo 40x/1.3 NA oil immersion objective. CACO-2_BBE_ cells were imaged using an Apo TIRF 100x/1.49 NA TIRF oil immersion objective. <u>Structured illumination microscopy</u> (SIM) was used for imaging frozen tissue sections and fixed cells with a Nikon N-SIM equipped with 405, 488, 468, and 647 nm LASERs, an Andor DU-897 EMCCD camera, and a TIRF 100X/1.49 NA TIRF oil immersion objective. All SIM images were reconstructed using Nikon Elements software.

### Live Imaging Microscopy

Prior to live cell imaging, cells growing in 35mm glass bottom dishes were rinsed once with 1X DPBS (Corning #21-031-CV). FluoroBrite imaging media (Gibco #A18967-01) supplemented with 10% FBS and 1% L-glutamine was added to the dish. For CL4 cells expressing Halo-CDHR2, Janelia Fluor 635 dye (Janelia) was added to the FluoroBrite media at a concentration of 50 nM for 1 hour at 37°C immediately prior to imaging. <u>Spinning disk confocal microscopy</u> was performed using a Nikon Ti2 inverted light microscope with a Yokogawa CSU-X1 spinning disk head, a Photometrics Prime 95B or Hamamatsu Fusion BT sCMOS camera, and three excitation LASERs (488, 568 and 647 nm). A 100X/1.49 NA TIRF oil immersion objective was used for all acquisitions. A stage incubator (Tokai Hit) maintained cells in a humidified environment at 37°C with 5% CO2.

### Fluorescence Recovery after Photobleaching (FRAP)

A square ROI was drawn in Nikon Elements at marginal and/or medial microvilli regions. A stimulating 405 nm LASER controlled by a Bruker mini-scanner set at 70% power and a dwell time of 40 us was targeted to each ROI after the first 3 frames of the movie acquisition. Two ND time acquisitions were used for imaging fluorescence recovery at intervals of 15 s for 3 minutes, followed by intervals of 30 s for 10 minutes.

### Electron Microscopy – CACO-2_BBE_ and LLC-PK1-CL4 cells and tissue

To prepare samples, cells were plated on glass coverslips washed once with warm SEM buffer (0.1M HEPES, pH 7.3) supplemented with 2 mM CaCl_2_, then fixed with 2.5% glutaraldehyde and then 4% paraformaldehyde in SEM buffer supplemented with 2mM CaCl_2_. Samples were washed in SEM buffer, then incubated in 1% tannic acid, washed with ddH_2_O, incubated with 1% OsO4, washed with ddH_2_O, incubated with 1% uranyl acetate, then washed with ddH_2_O. Samples were dehydrated in a graded ethanol series. Sampels were then dried using critical point drying and mounted on aluminum stubs and coated with gold/palladium using a sputter coater. SEM imaging was performed using Quanta 250 Environmental-SEM operated in high vacuum mode with an accelerating voltage of 5–10 kV, or imaged on a Zeiss Crossbeam 550 at 2keV. All reagents were purchased from Electron Microscopy Sciences. For more detailed methods, see [28]. TEM of mouse intestine (Fig. S1B) was performed as previously described [14]

### Electron Microscopy – Crypt-villus axis

For SEM imaging of intestinal sections, immediately after euthanasia, ∼5 mm murine duodenal sections were quickly fixed in a large volume (10mL) of 2.5% glutaraldehyde and 4% paraformaldehyde in SEM buffer (described above). Sections were then washed in SEM buffer prior to embedding in Tissue-Tek OCT compound (Sakura Finetek #4583). To ensure stable support of the complex architecture within the explant lumens, samples were gently moved through 3 rounds of fresh OCT compound with gentle manipulation to ensure penetration of the OCT. Samples were then placed in cryomolds (with OCT) and frozen over a dry ice/ethanol slurry. Molds were stored at -80C once fully frozen. Frozen explants were subsequently sectioned on a Leica CM1950 crytostat at 50μm/section and melted onto stainless steel AFM specimen discs (Electron Microscopy Sciences). Next, explant sections and disks were immersed in 1% OsO_4_, washed in ddH_2_O, then dehydrated through graded ethanol series. Of note, it was most common to experience detachment of the section from the AFM disk during the OsO_4_ and dehydration steps. Detached sections were recovered and gently adhered to an aluminum SEM specimen stub via conductive adhesive tab. SEM imaging was performed on a Quanta 250 environmental SEM, as described above.

### Quantification and Statistical Analysis

#### Microvilli Orientation Measurements

In Fiji, the first frames of three independent mCherry-ESPN CL4 cell movies were used for orientation measurements shown in Fig. 2B. A thin, rectangular ROI (height 12 pixels) was drawn across 2+ cells to encompass both marginal and medial areas (sample ROI Fig 2A, dotted box). The ROI hyperstack was duplicated and 3D projected with rotation around the X axis. Using the Angle tool, a line was drawn down the length of each microvillus (dotted lines, Fig. 2B) with the angle base parallel to the cell surface (solid lines Fig. 2B). Angle measurements were plotted in Prism in a column chart and mean marginal and medial angles were compared using a Welch’s unpaired t-test.

#### Temporal Color Coding

Time frames for every 3 minutes were selected (18 total frames). Using the Temporal-Color Code function in Fiji, the ESPN channel was coded (start frame 1, end frame 18) using the Spectrum LUT (Fig. 3B).

#### Microvilli tracking using EGFP-EPS8 puncta

Denoised and deconvolved 3D movies were converted into max intensity projections in the Z plane. Next, a binary via the spot detection tool was applied to the FITC channel (EPS8 signal) with a diameter of 260 nm and a contrast value of 25 to threshold EPS8 puncta representing individual microvilli in the ROI (medial or marginal). Tracking parameters did not allow for the detection of new tracks after the first frame, allowed for a maximum of 3 gaps in a given track, and a standard deviation multiplier of 2. Using the tracking tool, binaries, representing EPS8 puncta, were tracked and any points lying outside of the ROI were deselected. Track data, time and X Y positions, were then exported to Excel and analyzed and plotted in Prism as radial X Y positions over time by subtracting each position in X or Y from the respective point position at time 0, making the first position (0,0) (Fig 3C and 3F). Three independent live cell imaging experiments were used for the analysis.

#### Mean Square Displacement

With the same X,Y EPS8 puncta trajectories described above, an Excel spreadsheet was used to calculate the mean square displacement of each tracked microvillus over 15 s intervals across 5 minutes as us and others have done previously [28, 49–51]. A line was fitted to each MSD curve using the simple linear regression model in Prism which provided the slopes presented on the plots in Fig. 3E and H.

#### Cell Mixing Linescans

Using Fiji, a segmented line with a width of 6 was drawn across mixed cell interfaces to encapsulate signal at the mixed cell (marginal) interfaces. Lines with a minimum length of 20 µm and maximum length of 75 µm were used in analysis. Fig. 5 shows one representative linescan from the large dataset for each cell mixing scenario, with line length (µm) shown on the X axis and mCherry and EGFP construct intensities on the Y axis. Intensities were normalized from 0 to 1 in Prism with 0 being the lowest gray value in the linescan and 1 being the highest. The residual plots shown were calculated from the respective representative linescan by subtracting mCherry intensity from EGFP intensity at each length in X. 30 individual linescans from each cell mixing scenario were plotted on their own XY correlation plot in Prism. Combined Pearson’s r values from the 30 individual correlations were plotted in Fig 5K, and mean r values were compared in Prism using an Ordinary one-way ANOVA with multiple comparisons.

#### FRAP Fraction Recovery Analysis

A background ROI and reference ROI were used to account for photobleaching and background fluorescence in both channels. Fraction recovery over time was calculated from 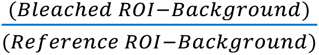. Recovery curves were fitted with a two-phase association equation in Prism and the immobile fraction was calculated from 1 minus the plateau. Images shown in Fig. 6 were denoised and deconvolved in Nikon Elements for presentation clarity, however all analyzed measurements presented in the FRAP plots were taken from raw, unprocessed movies.

#### Mean ESPN Intensity Measurements

In Nikon Elements, the movie was projected in Z to create a maximum intensity projection. For Fig. 7D, at 0 hr and 24 hr a ring ROI was drawn encompassing each marginal zone and separate circular ROI was drawn to encompass the medial zone of 10 cells. Mean mCherry-ESPN intensity was measured for each cell at the marginal and medial zones. Delta represents the change in mCherry-ESPN mean intensity from the 0 hr time point to the 24 hr time point.

## SUPPLEMENTAL FIGURE LEGENDS

**Figure S1:**
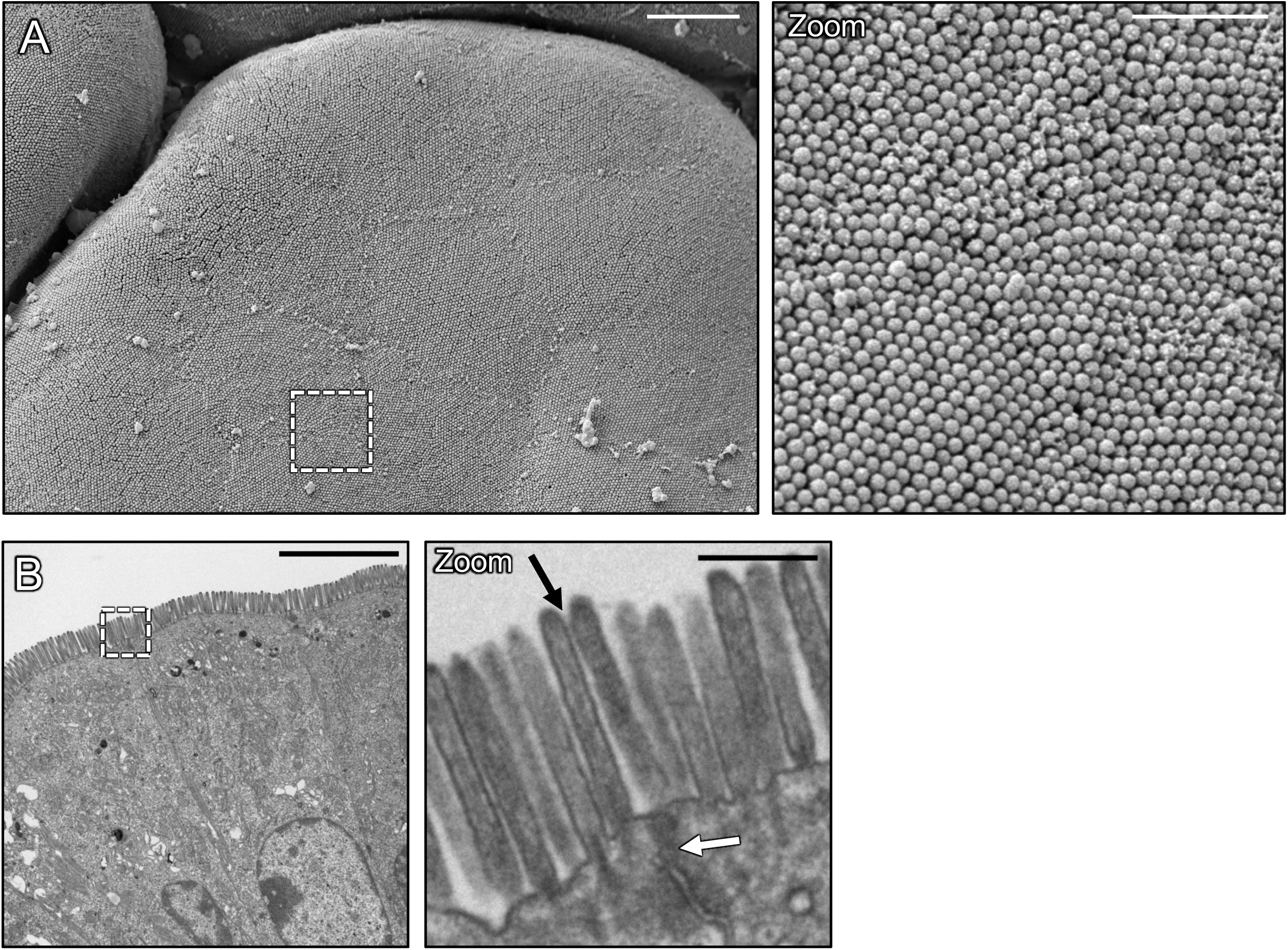
Microvilli of mature epithelial cells are continuous across the monolayer surface. (**A**) SEM of mouse small intestine showing an *en face* view of microvilli on neighboring enterocytes. The dashed box represents a zoom of a single enterocyte cell and its neighbors. (**B**) Transmission electron micrograph (TEM) of mouse small intestine showing a lateral view of microvilli. Dashed box represents the zoom of a cell-cell junction (white arrow) with neighboring cell microvilli appearing continuous across the interface (black arrow). Scale bars: 4 µm (A), 1 µm (A, zoom), 5 µm (B), 500 nm (B, zoom).

**Figure S2:**
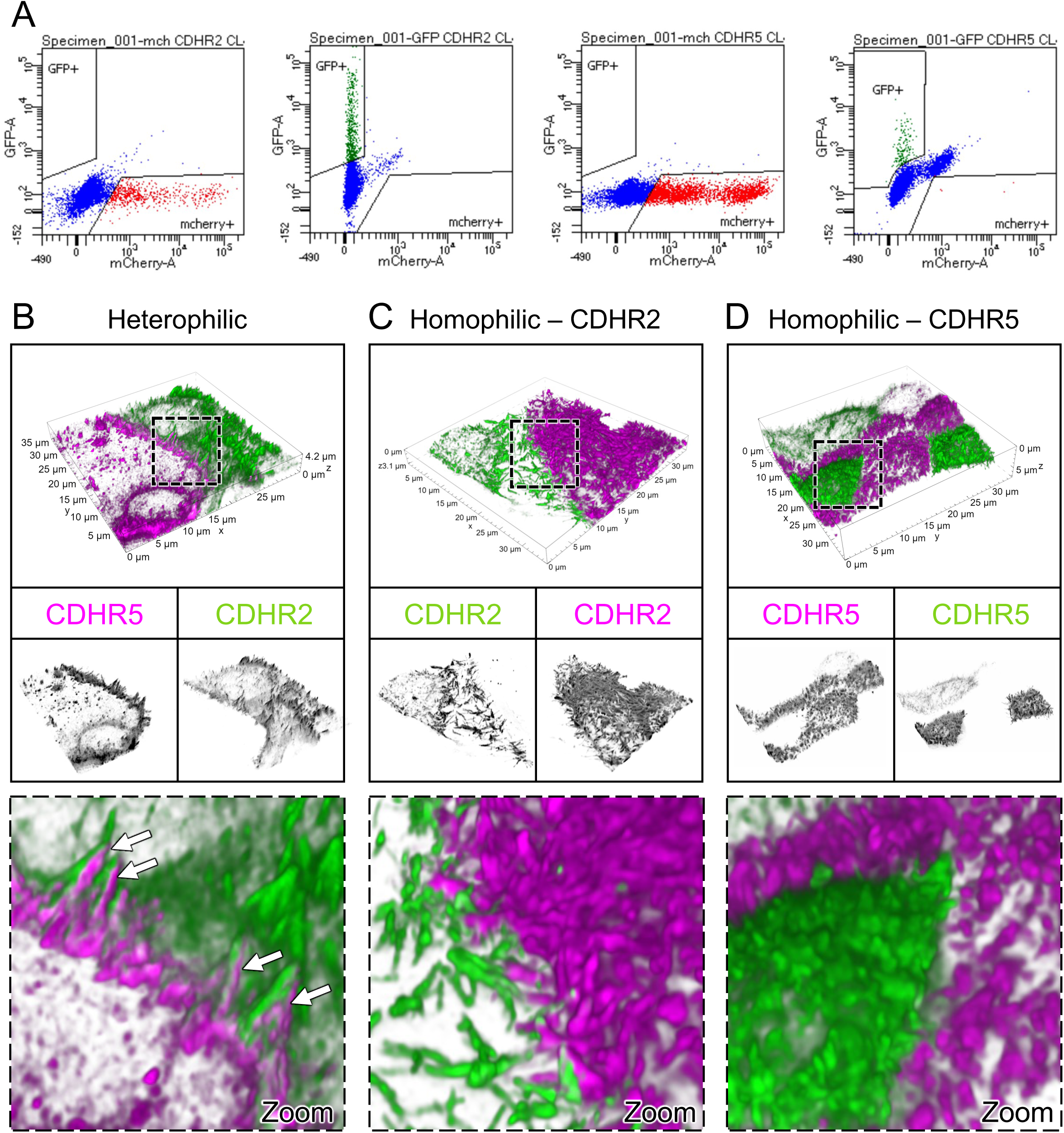
High-resolution imaging of adhesion complex interfaces in mixed CL4 cell populations. **(A)** FACS profiles of the four stable CL4 cell lines expressing C-terminal mCherry or EGFP tagged CDHR2 or CDHR5 as marked on the top axis of the graph. 3D Volume SIM images of mixed **(B)** heterophilic; CDHR5-mCherry and CDHR2-EGFP, **(C)** homophilic; CDHR2-EGFP and mCherry-CDHR5, and **(D)** homophilic; CDHR5-mCherry and CDHR5-EGFP CL4 cells. Dashed boxes outlined in B-D represent zooms shown in bottom panel, respectfully. White arrows in zoom under (B) point to instances of robust microvilli alignment at cell margins, which are absent in the homophilic mixing scenarios. Scales as marked.

## SUPPLEMENTAL VIDEO LEGENDS

**Video S1: Microvilli adopt an orientation perpendicular to the cell surface upon reaching the cell margins.** 3D volume projection depth coded in Z of live mCherry-ESPN expressing CL4 cells. Spinning disk confocal movie taken over 2 hours with 2-minute intervals of the microvilli cluster marked in Fig 2C, and D. Arrow follows a cluster of medial microvilli that transition to a vertical orientation upon reaching the marginal cell area (as marked by Z-depth color profile). Scale bar: 3 µm.

**Video S2: Marginal heterophilic adhesion complexes have low signal recovery, suggesting high complex stability.** Mixed CL4 cells expressing CDHR2-EGFP (green) and CDHR5-mCherry (magenta) photobleached at two marginal ROIs; 30s intervals shown. Scale bar: 5 µm.

**Video S3: Medial heterophilic adhesion complexes have higher signal recovery, suggesting lower complex stability.** CL4 cell co-expressing CDHR2-Halo (magenta) and CDHR5-EGFP (green) photobleached at a medial ROI; 15s and 30s intervals shown. Scale bar: 5 µm.

**Video S4: Marginal homophilic CDHR2 adhesion complexes have higher signal recovery, suggesting lower complex stability.** Mixed CL4 cells expressing CDHR2-EGFP (green) and CDHR2-mCherry (magenta) photobleached at a marginal ROI; 15s intervals shown. Scale bar: 5 µm.

**Video S5: Marginal homophilic CDHR5 adhesion complexes have higher signal recovery, suggesting lower complex stability.** Mixed CL4 cells expressing CDHR5-EGFP (green) and CDHR5-mCherry (magenta) photobleached at a marginal ROI; 30s intervals shown. Scale bar: 5 µm.

**Video S6: Microvilli accumulate first at cell margins over cell surface differentiation.** (Left) CL4 cells expressing mCherry-ESPN imaged at 3-minute intervals for 43 hours. (Right) Fire LUT intensity profile of the ESPN channel. Intensity scales from low (0; dark purple) to high (255; yellow/white) as denoted by LUT profile. StackReg ImageJ plugin was used to align all frames of the movie. Scale bar: 10 µm.

