## Supplemental Figures 1 and 2 for "Adhesion-based capture stabilizes nascent microvilli at epithelial cell junctions"

#### SUPPLEMENTAL FIGURE LEGENDS

**Figure S1: Microvilli of mature epithelial cells are continuous across the monolayer surface.** (**A**) SEM of mouse small intestine showing an *en face* view of microvilli on neighboring enterocytes. The dashed box represents a zoom of a single enterocyte cell and its neighbors. (**B**) Transmission electron micrograph (TEM) of mouse small intestine showing a lateral view of microvilli. Dashed box represents the zoom of a cell-cell junction (white arrow) with neighboring cell microvilli appearing continuous across the interface (black arrow). Scale bars: 4 μm (A), 1 μm (A, zoom), 5 μm (B), 500 nm (B, zoom).

**Figure S2: High-resolution imaging of adhesion complex interfaces in mixed CL4 cell populations. (A)** FACS profiles of the four stable CL4 cell lines expressing C-terminal mCherry or EGFP tagged CDHR2 or CDHR5 as marked on the top axis of the graph. 3D Volume SIM images of mixed **(B)** heterophilic; CDHR5-mCherry and CDHR2-EGFP, **(C)** homophilic; CDHR2-EGFP and mCherry-CDHR5, and **(D)** homophilic; CDHR5-mCherry and CDHR5-EGFP CL4 cells. Dashed boxes outlined in B-D represent zooms shown in bottom panel, respectfully. White arrows in zoom under (B) point to instances of robust microvilli alignment at cell margins, which are absent in the homophilic mixing scenarios. Scales as marked.

### Fig. S1

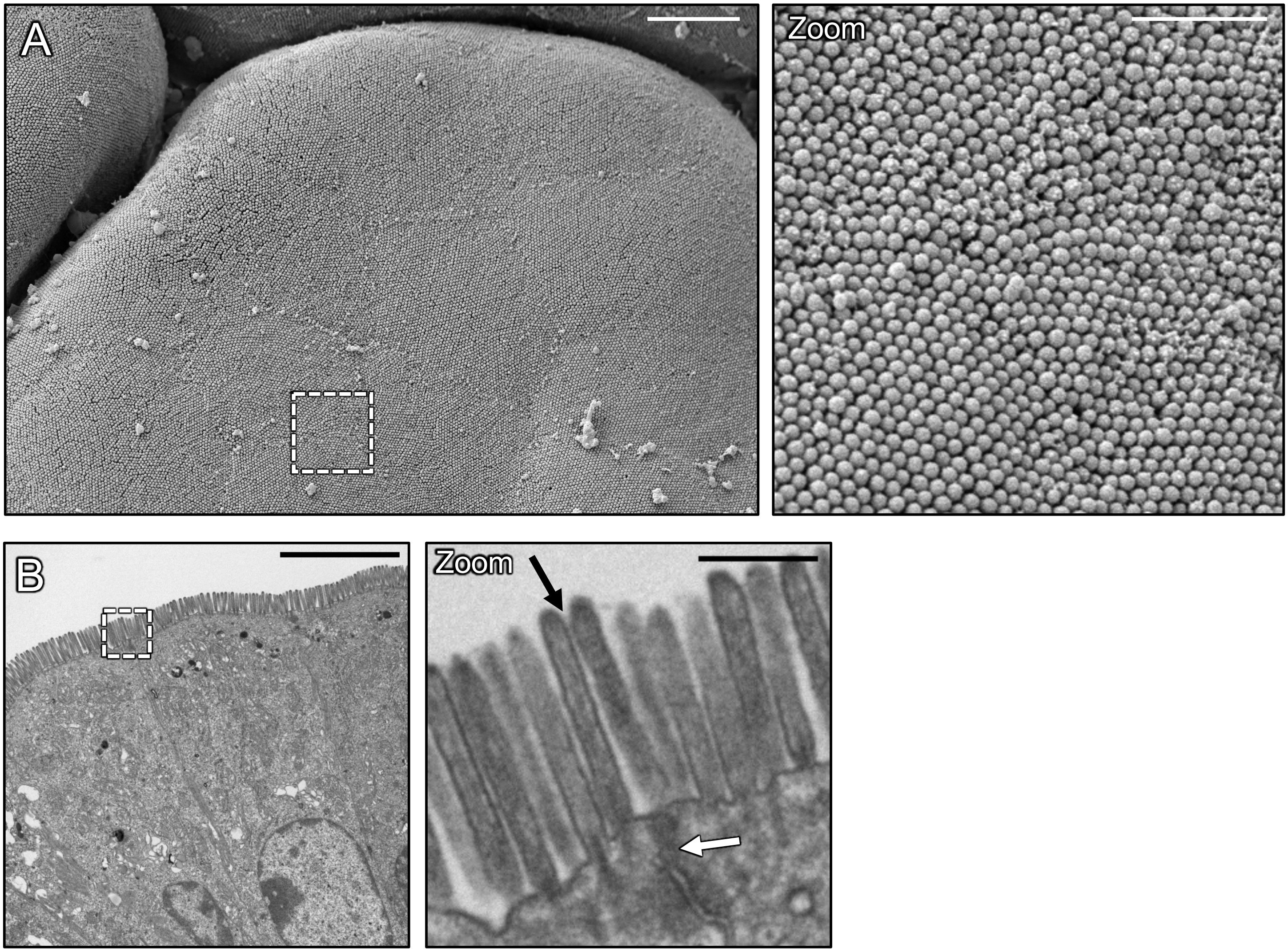

### Fig. S2

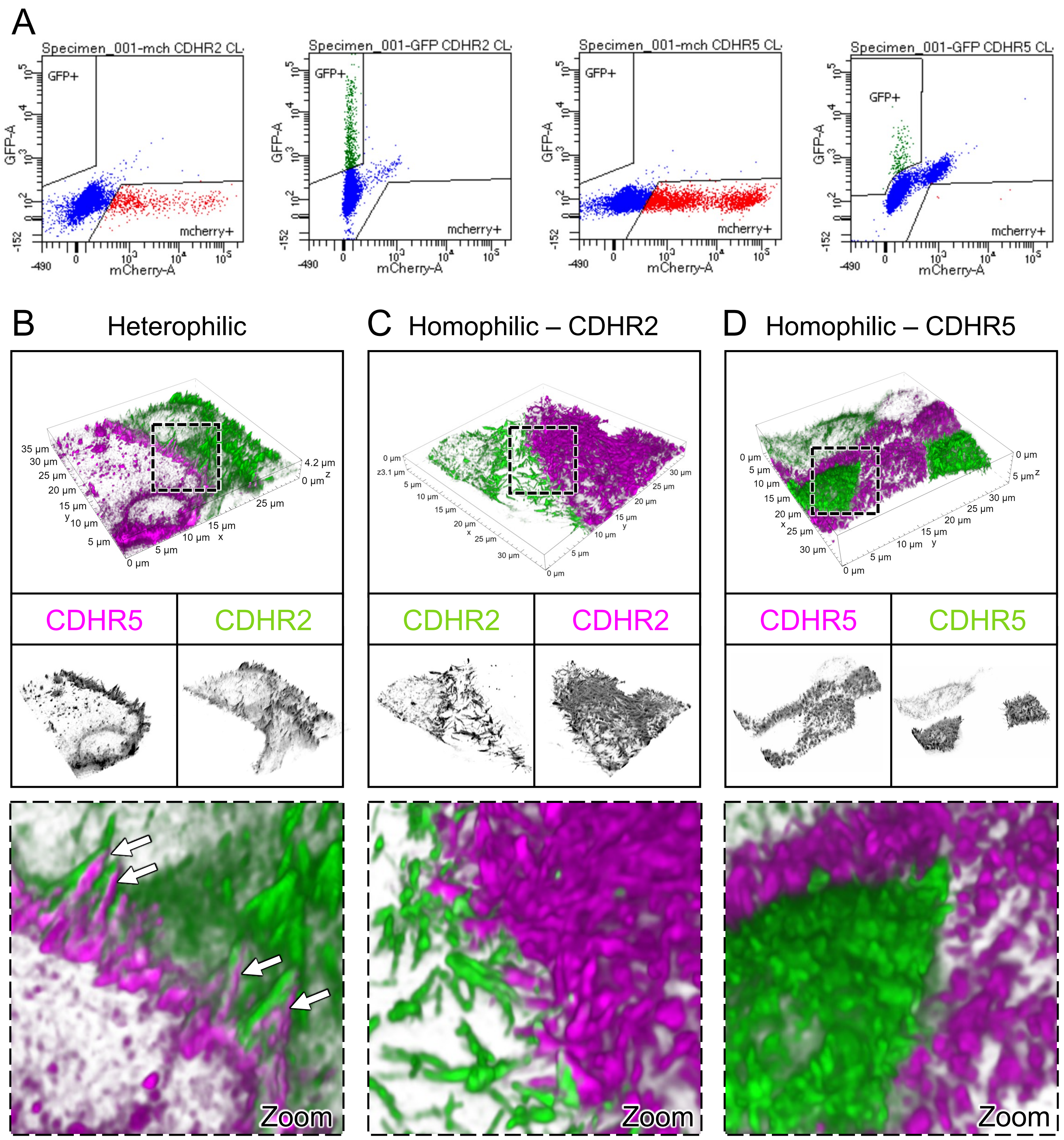
A
Specimen_001-mch CDHR2 CL
Specimen_001-GFP CDHR2 CL
Specimen_001-mch CDHR5 CL
Specimen_001-GFP CDHR5 CL
GFP+
mcherry+
GFP-A
mCherry-A
B
Heterophilic
C
Homophilic – CDHR2
D
Homophilic – CDHR5
CDHR5
CDHR2
CDHR2
CDHR2
CDHR5
CDHR5
Zoom
Zoom
Zoom
